# Secondary Structure Motifs Made Searchable to Facilitate the Functional Peptide Design

**DOI:** 10.1101/651315

**Authors:** Cheng-Yu Tsai, Emmanuel O Salawu, Hongchun Li, Guan-Yu Lin, Ting-Yu Kuo, Liyin Voon, Adarsh Sharma, Kai-Di Hu, Yi-Yun Cheng, Sobha Sahoo, Lutimba Stuart, Chih-Wei Chen, Yuan-Yu Chang, Yu-Lin Lu, Ximai Ke, Chen-Chi Wu, Chung-Yu Lan, Hua-Wen Fu, Lee-Wei Yang

## Abstract

To ensure a physicochemically desired sequence motif to adapt a specific type of secondary structures, we compile an α-helix database allowing complicate search patterns to facilitate a data-driven design of therapeutic peptides. Nearly 1.7 million helical peptides in >130 thousand proteins are extracted along with their interacting partners from the protein data bank (PDB). The sequences of the peptides are indexed with patterns and gaps and deposited in our Therapeutic Peptide Design dataBase (TP-DB). We here demonstrate its utility in three medicinal design cases. By our pattern-based search engine but not PHI-BLAST, we can identify a pathogenic protein, *Helicobacter pylori* neutrophil-activating protein (HP-NAP), a virulence factor of *H. pylori*, which contains a motif DYKYLE that belongs to the affinity determinant motif DYKXX[DE] of the FLAG-tag and can be recognized by the anti-FLAG M2 antibody. By doing so, the known purification-tag-specific antibody is repurposed into a diagnostic kit for *H. pylori*. Also by leveraging TP-DB, we discovered a stretch of helical peptide matching the potent membrane-insertion pattern WXXWXXW, elucidated by MD simulations. The newly synthesized peptide has a better minimal inhibitory concentration (MIC) and much lower cytotoxicity against *Candida albicans* (fungus) than that of previously characterized homologous antimicrobial peptides. In a similar vein, taking the discontinued anchoring residues in the helix-helix interaction interface as the search pattern, TP-DB returns several helical peptides as potential tumor suppressors of hepatocellular carcinoma (HCC) whose helicity and binding affinity were examined by MD simulations. Taken together, we believe that TP-DB and its pattern-based search engine provide a new opportunity for a (secondary-)structure-based design of peptide drugs and diagnostic kits for pathogens without inferring evolutionary homology between sequences sharing the same pattern. TP-DB is made available at http://dyn.life.nthu.edu.tw/design/.

## 1. Introduction

Secondary structures comprise a subset of amino acid sequences where each constituent residue can form hydrogen bonds (H-bonds) in specific orders with other constituent residues, at their peptidyl backbones, which grants a special mechanical property in our life-thriving nanomachines – proteins. Particularly, α-helices, where the *i*-th residue in the helix H-bonded with *i*+4-th residue, mediate various types of molecular functions including cell penetration^1, 2, 3, 4^, subcellular localization^5^, intra/inter-molecular allostery^6–9^, special helix-DNA scaffolds^10^ and protein-protein interaction^7, 11^, provided that the primary sequence meets both the structural (main chain and side chain) and chemical (side chain) requirements for specific functions of interest. Especially at the interface, either water-lipid interface at the cell membrane, or protein-DNA/protein-protein interface, these helices have disparate physicochemical properties at their two sides to properly play their functional roles. Take amphiphilic antimicrobial peptides (AMPs) for example, if the *i*-th position is a membrane-insertion-promoting amino acid (say, W or Y), the *i*+3 or *i*+4-th position needs to be an amino acid also prompt for membrane insertion (say, another hydrophobic residues). In other words, these residues with similar physicochemical properties need to be aligned at the same side, while the other side faces bulk water and ions accumulated near the membrane. In this example, what we need is a stretch of amino acid sequence consisted of a pattern of (W/Y)X_1_X_2_(W/Y) or (W/Y)X_1_X_2_X_3_(W/Y), where X_i_ can be any amino acid as long as the combination of Xi can still hold an helical structure, and (W/Y) means that position can be either W or Y. From the point of view of protein design, either to design a better AMP or to propose stronger interface binders, and therefore blockers, for therapeutic purposes, it is not an easy task to allow constituent residues to address physicochemical needs, meeting a specific sequence pattern, while maintaining the required structural integrity. Suitable tools need to be developed to facilitate the process.

Currently, both (A) pattern-based (only) search engines locating desired functional motifs in primary sequences *regardless of their evolutionary conservation* and (B) dedicated secondary structural databases indexed for (A)’s purpose are lacking. Protein Secondary Structure database (PSS) proposed 30 years ago no longer serves the community^12^, while DSSP annotates the secondary structures when provided with tertiary structures of proteins^13^. However, both DSSP and protein databank (PDB) do not provide a search engine for specific types of secondary structures (say, the entire query sequence should belong to an alpha-helix) for >170,000 structurally solved sequences, nor do they provide any search function that recognizes specific sequence patterns (see the beginning of Methods for our definition of “patterns”). To better integrate functional structural fragments into therapeutics^14, 15, 16^, in this seminal work, we extracted ∼1.7 million (1,676,117) structurally resolved helices and their contacting/interacting helical partners from the PDB, from which helical propensity and coordination (contact) number are derived (Table 1). We then establish a pattern-specific search engine (see below) to find indexed sequences bearing particular spatio-chemical properties instead of evolutionary homology (say, the purpose of Pattern Hit Initiated BLAST (PHI-BLAST)^17^ that takes input of both residue pattern and the template sequence). We use this Therapeutic Peptide Design dataBase (TP-DB) to develop a new antifungal/antimicrobial peptide, helix-helix interface blockers that suppress hepatocellular carcinoma, and a new diagnostic reporter to detect *Helicobacter pylori* infection.

**Table 1.** Amino acids sorted by their indexes of helical propensity

| AA* | HP<sub>a.a.</sub> | NA<sub>a.a.</sub> | HP<sub>NA</sub><br>(HP<sub>a.a.</sub>/NA<sub>a.a.</sub>) | Helical Propensity,<br>log (HP<sub>NA</sub>) |
| --- | --- | --- | --- | --- |
| A | 0.109 | 0.083 | 1.320 | 0.277 |
| W | 0.014 | 0.011 | 1.284 | 0.250 |
| E | 0.084 | 0.067 | 1.246 | 0.220 |
| L | 0.115 | 0.097 | 1.192 | 0.175 |
| Q | 0.045 | 0.039 | 1.145 | 0.135 |
| Y | 0.033 | 0.029 | 1.130 | 0.122 |
| K | 0.064 | 0.058 | 1.100 | 0.095 |
| M | 0.026 | 0.024 | 1.079 | 0.076 |
| R | 0.059 | 0.055 | 1.067 | 0.065 |
| F | 0.038 | 0.039 | 0.984 | -0.016 |
| I | 0.058 | 0.059 | 0.978 | -0.022 |
| H | 0.022 | 0.023 | 0.969 | -0.031 |
| D | 0.052 | 0.055 | 0.952 | -0.049 |
| N | 0.038 | 0.041 | 0.936 | -0.066 |
| T | 0.049 | 0.053 | 0.918 | -0.086 |
| V | 0.062 | 0.069 | 0.902 | -0.103 |
| C | 0.012 | 0.014 | 0.876 | -0.132 |
| S | 0.053 | 0.066 | 0.804 | -0.218 |
| G | 0.046 | 0.071 | 0.650 | -0.431 |
| P | 0.021 | 0.047 | 0.445 | -0.810 |

\* AA = Amino Acid; HP<sub>a.a.</sub> = Helical Propensity for each amino acid; NA<sub>a.a.</sub> = Natural Abundance of each amino acid; HP<sub>NA</sub> = HP<sub>a.a.</sub> normalized by NA<sub>a.a.</sub>

## 2. Methods

### 2.1. Extraction of Helical Peptide Sequences from the Protein Data Bank (PDB)

To obtain the amino acid sequences that fold into helices in nature, we processed the PDB^18^ files of 130,000+ experimentally determined protein structures as of June, 2017 and extracted the locations of secondary structures from the header part of the PDB files. There are 96,711 unique PDB IDs/files containing at least one alpha-helix documented at the header area to result in 1,676,087 structurally resolved helices having a length longer than 5 residues. In addition, for a given helix, its contacting neighbors in 3D space and the adjacent helical peptides that are within 4Å (per heavy atoms) were also collected and analyzed.

### 2.2. Indexing and Querying of TP-DB

Helical peptides that match specific patterns are carefully indexed in the database. A pattern could be of the form “Y***G**K”, where “*” belongs to any of the 20 amino acids, equivalent to the regular expression “Y-x(3)-G-x(2)-K” in a PROSITE format that has also been used in the PHI-BLAST search (https://www.ncbi.nlm.nih.gov/blast/html/PHIsyntax.html). The location of a pattern is indexed by the PDB ID, the chain ID, and the position where the pattern begins in that chain (see below). Considering both flexibility and search comprehension, we let the simplest pattern be [A_1_ m A_2_ n A_3_] where A_1_, A_2_, and A_3_ are the single-letter codes of certain amino acids, in between which are the spacings represented by the whole numbers m and n. We currently allow m and n to be from zero to four. A key “Y3G2K” or its search pattern [Y 3 G 2 K] would indicate a “Y” is (3 + 1) residues upstream to a “G” that is (2 + 1) upstream to a “K”. A key is paired with identifiers (values) referring to where this key can be found in all the stored peptides (Fig. 1C, E). Specifically, these values, stored in an array, are peptide identifiers and all the starting positions of a certain key in each of the peptides (**Fig. 1C**). These key-value pairs comprise our database (**Fig. 1**), TP-DB (http://dyn.life.nthu.edu.tw/design/), which could be searched by a simple pattern [A_1_ m A_2_ n A_3_] or a pattern compounded by several of these simple patterns (**Fig. 1F**).

**Fig. 1.**
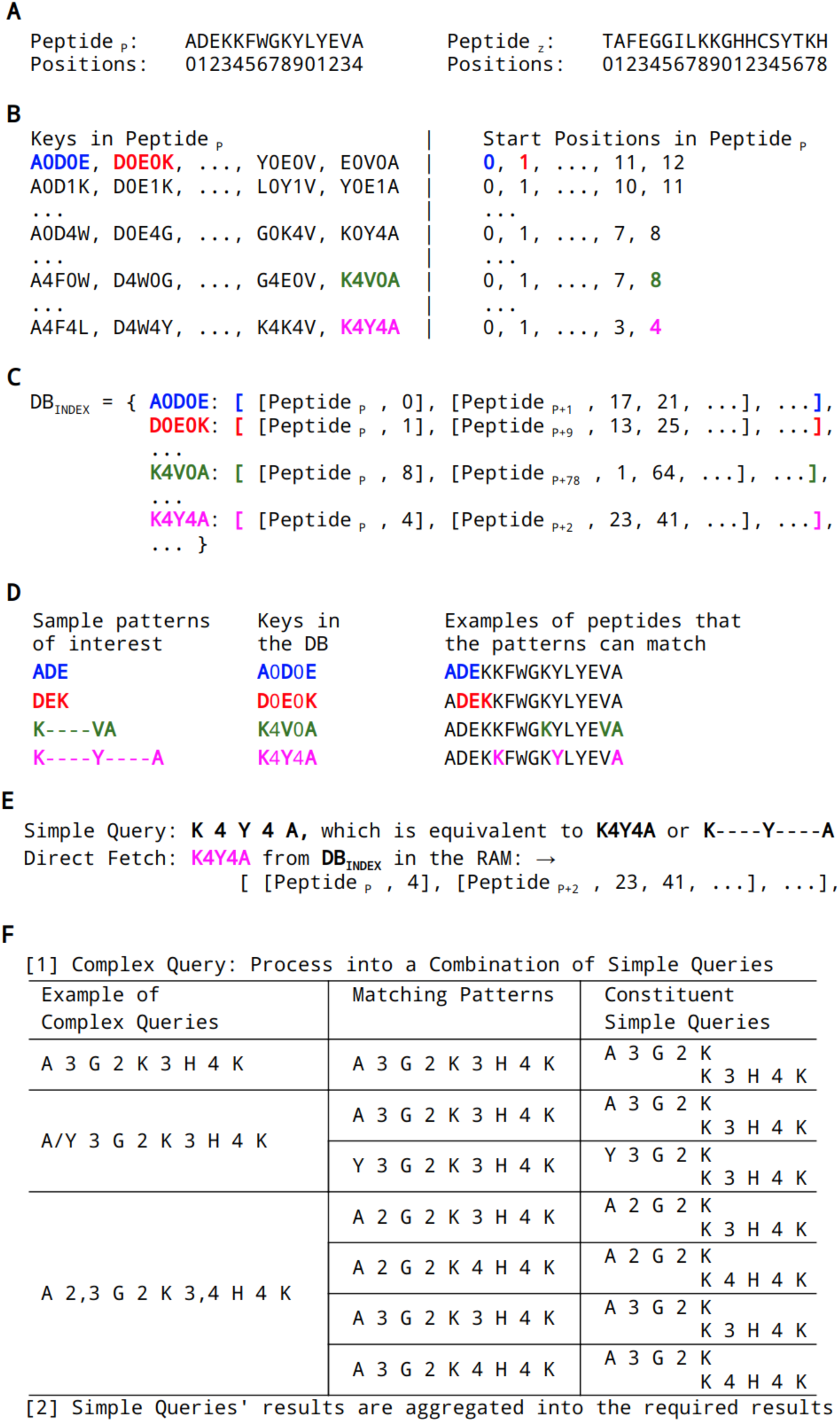
Indexing the peptides and querying of TP-DB. (**A**) The positions of amino acids in the *p*-th helical peptide (peptide_p_) are indexed from 0 to 14. (**B**) All the possible patterns that can be found in peptide_p_, stored as keys and values. (**C**) In TP-DB (or DB_Index_), keys are paired values, and values are where these keys can be found, including in which peptides and the starting position of the keys. For instance, [A 0 D 0 E] can be found in the “0-th position” in the *p*-th peptide, while it can also be found in the 17th and 21st positions in the (*p*+1)-th peptide. (**D**) The patterns of interest are translated into “keys” comprising amino acids’ single-letter codes separated by numbers, the sizes of the gaps. (**E**) Results for a simple query retrieved from TP-DB. (**F**) A compounded query can be broken down into subqueries and the result is a joint result from subqueries. For efficient search, two types of libraries are loaded in timely order. What is constantly loaded in the memory, with a small memory footprint, is a collection of key-value pairs that only index which proteins have certain keys but not their locations. Only when certain proteins are visited because they contained the queried keys, then libraries of a second type, inferring the locations of keys in certain proteins, are then loaded into the memory to report the found sequences, before the protein-relevant libraries are unloaded from the memory again soon after a search job is finished.

To query the database, the user specifies at least three anchor residues and the gaps that separate them, such that “ADE” or “K****VA” can be queried by [A 0 D 0 E] and [K 4 V 0 A], respectively (**Fig. 1D**). For more complicated queries, say “A/Y 3 G 2 K 3 H 4 K”, it is first broken down into a combination of two subqueries “A 3 G 2 K 3 H 4 K” and “Y 3 G 2 K 3 H 4 K” (**Fig. 1F**). The five anchor residues in each subquery is treated as two concatenated three-anchor keys. The matched patterns co-localized in a peptide sequence are further parsed to take into account the overlapped regions before returning the results. TP-DB currently have two search engines to search the indexed peptides, one by pattern search, described above, and the other by BLAST (parameterized for short sequences). The use of the latter is beyond the scope of this paper.

### 2.3. Physicochemical Properties and Ranking of the Pattern-Matched Helical Peptides in TP-DB

The helical peptides returned by a query can be ranked by (the logarithm of) their normalized helical propensity (default) or by their contact scores. Using the data collected in the TP-DB, each of the 20 amino acids has a normalized helical propensity HPNA that is HP_a.a._/NA_a.a._ where HP_a.a._ is the helical propensity of a given amino acid type and NAa.a is natural abundance of that amino acid (**Table 1**). Both HP_a.a._ and NA_a.a._ are probabilities, where HP_a.a_ is defined as the count of a certain amino acid type in all the extracted helices divided by the total number of amino acids in these helices (24,301,682 amino acids in 1,676,117 helical peptides), and NA_a.a._ is the count of that amino acid in the 130,000+ proteins in PDB, divided by the total number of residues in these proteins. A log-odds ratio, log (HPNA) or log (HP_a.a._/NA_a.a._) can be found for each amino acid type, and the reported helical propensity for each peptide in TP-DB is the sum of log (HPNA) for every constituent residue in this peptide (**Table 2**). The contact number is the amino acid contact within a 15Å range of a residue of interest, in the residue’s original protein environment, where only the Cα atom of an amino acid is considered. The reported contact for a peptide in TP-DB is the per-residue average of its constituent residues’. An exemplified output table from the web server is shown in **Table 2**.

**Table 2.**
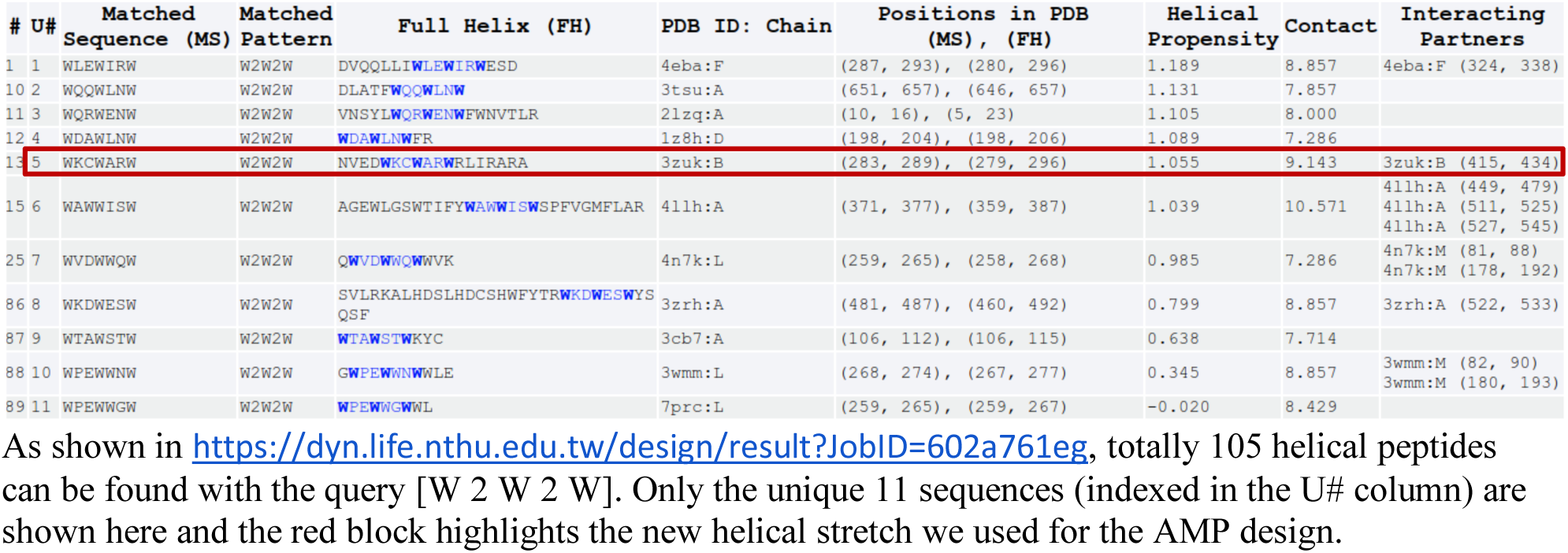
Results obtained from the TP-DB for the query [W 2 W 2 W]

### 2.4. General MD Simulations Protocol

All MD simulations were performed using NAMD package 2.9^19^ and OpenMM^20^ in NPT ensemble with periodic boundary conditions. Latest CHARMM forcefields^21, 22^ were employed for the peptide, water and lipids, respectively. All simulations were performed at a time step of 2 fs and trajectories were recorded every 5000 steps (i.e. at every 10 ps). RATTLE and SETTLE algorithms are applied to constrain hydrogen atoms in peptides and waters. Cutoff of 12 Å with switch distance 10 Å and pair list distance 14 Å are applied when calculating non-bonded interactions. With periodic boundary conditions, the Particle Mesh Ewald method^23^ was employed for calculations of long-range electrostatic interactions. Temperature was maintained at 310 K using Langevin dynamics^24^ and pressure was controlled at 1 atm using Nosé-Hoover Langevin piston^25^.

### 2.5. Lipid Preparation and Equilibrium *in silico*

Zwitterionic membranes with 78 1-palmitoyl-2-oleoyl-sn-glycero-3-phosphocholine (POPC; also termed as “PC” hereafter) lipids to mimic eukaryotic membrane and with 30 POPC lipid + 10 2-oleoyl-1-pamlitoyl-sn-glyecro-3-glycerol (POPG; also termed as “PG”) lipid molecules (see **Fig S1**) in each of the upper and lower lipid bilayers to mimic pathogenic prokaryotic membrane^26, 27^ were prepared by CHARMM-GUI server^28^. The AMPs and lipid bilayers were solvated in explicit TIP3 water molecules^29^. Na^+^ and Cl^-^ ions were added to neutralize the system. The initial area per lipid of PC was equilibrated to stay close to its experimental values measured at 303 K, which is 68.3 ± 1.5 Å^2 30^.

### 2.6. Natural insertion simulations with soft-boundary-confined diffusion for AMPs in the presence or absence of lipids

NMR-solved α-helical and fully extended (random-coil) AMPs were simulated together with PC and PC/PG (3:1) membranes in a water box containing 0.02 M NaCl. For the fully extended AMPs, they were first simulated in an NPT ensemble for 50 ns in the absence of lipids. The resulting snapshots of the AMPs were clustered into 5 groups based on their structural similarity using the “clustering” plug-in of the VMD software^31^. A representative conformation from the biggest cluster was selected as the initial conformation for an AMP before it was loaded with the equilibrated membrane for the natural insertion simulations. The expression “natural” meant that no external pulling forces were exerted on the AMPs, contrasting to an accelerated approach we developed earlier^2^ to first pull the AMPs into the lipids and release them in order to timely assess the insertion quality of AMPs.

### 2.7. Determination of the Minimum Inhibitory Concentration (MIC)

To measure antimicrobial activity of peptides, minimum inhibitory concentrations (MIC) were determined by broth-dilution assay as described in Clinical and Laboratory Standards Institute (CLSI) documents M27-A3 for yeasts^32^ and M07-A9 for bacteria^33^ with minor modifications. Briefly, *E. coli* ATCC 25922 and *C. albicans* SC5314 were recovered from frozen stock by growing overnight at 37°C on LB (Luria-Bertani) and YPD (Yeast Extract-Peptone-Dextrose) agar plates, separately. A single colony of *E*. *coli* was inoculated into 5 ml LB medium and grown overnight at 37°C with shaking at 220 rpm. Cells were then diluted into MHB (Mueller–Hinton broth) at 1:100, and grew until the optical density at 600 nm (OD_600_) reached 0.4 to 0.8. For *C. albicans* SC5314, a single colony was inoculated in YPD broth and grown overnight at 30℃ with shaking at 180 rpm. Cells from the overnight culture were subcultured into fresh YPD and grew to mid-log phase. Finally, *E*. *coli* and *C*. *albicans* cells obtained from above-described were diluted into MHB and LYM, a modified RPMI 1640 medium^34^ to a concentration of ∼5 × 10^5^ cells/ml and ∼4 × 10^4^ cells/ml, respectively. To determine the MIC values, one hundred microliters of cells was placed in each well of a clear bottom 96-well microplate, followed by independently adding 100 µl peptides with different concentrations (in MHB or LYM). The final concentrations in the wells were 120, 60, 30, 15, 7.5, 3.75 and 1.88 µg/ml for peptides, ∼ 2.5 × 10^5^ cells/ml for *E. coli* and ∼2 × 10^4^ cells/ml for *C. albicans*. The plate was incubated at 37°C with shaking at 220 rpm for 18 h (*E. coli*) and at 180 rpm for 24 h (*C. albicans*). The OD_600_ values were measured using a iMark^TM^ Microplate Absorbance Reader (Bio-Rad). The MIC_90_ value was determined by 90% reduction in growth compared with that of the peptide-free growth control. Experiments were performed in pentaplicate, and MIC_90_ was determined as the majority value out of 5 repeats.

### 2.8. Assays for Minimum Hemolytic Concentration (MHC)

The hemolytic toxicity was determined by the hemolysis against human red blood cells (RBCs) as previous publication^35^. First, RBCs from healthy donors were mixed with EDTA in the collecting tubes to inhibit the coagulation. Then the RBCs were washed for three repeats as follows - the centrifugation at RCF 800g for 10 minutes, the removement of the supernatant, and the slight resuspension with PBS buffer. Such washed RBCs stock was finally diluted with a PBS buffer to be the 10% (v/v) RBCs solution for assays. Then total eight samples of various final concentrations (240, 120, 60, 30, 15, 7.5, 3.75 and 1.88 µg/ml) for each AMP in 96-well microplate were prepared by mixing 10% RBCs and diluted AMP solutions, respectively, at ratio 1:1 (v/v) into final volume 200 µL. All samples were placed under the condition of 37°C for 1 h and then centrifuged at RCF 800g for 10 minutes. Finally, all these supernatants from samples and controls were picked out for the measurement of optical density at the 450nm wavelength. The data processing was performed to calculate the hemolytic percentage defined as [A_sample_ - A_control_-]/[A_control_+ - A_control_-], where A_control_+ and A_control_- are the measurement at wavelength 450nm for the positive and negative controls, that are 100 µL 10% RBCs suspended in 100 µL PBS buffer with and without 2% (v/v) Triton X-100 (a known detergent to lyse the RBCs), respectively. The final hemolytic percentage for AMPs at various concentrations were determined by the mean values of three repeated experiments, and MHC5 values that the minimal concentration of AMPs leading to 5% hemolysis of RBCs were determined. The protocol was pre-approved by Research Ethics Committee of the National Taiwan University Hospital (approval number: 201810004RINA), and blood collections were performed by trained phlebotomists to confirm the safety of the donors.

## 3. Results and Discussions

### 3.1. Database Features and Statistics

TP currently comprises 1,676,117 helical peptides (24,301,682 amino acids in total) extracted from 130,000+ experimentally solved protein structures stored in PDB. The apparent goal of establishing search engines for protein secondary structures (starting from all types of helical configurations, e.g., α-helices and 310 helices.) is to find stretches of protein sequences that match the queried pattern and adapt a helical conformation. Since protein sequences assume structures in the context of their 3-dimensional environment full of residue contacts, it is quite likely the helices are not as helical as when they are part of their parent proteins. We herein design two metrics to evaluate and prioritize the peptides, in the query results, which are highly likely to still stay helical when expressed or synthesized alone - helical propensity (HP) and contact number. A higher HP and/or lower contact number would suggest a good helicity for peptides even in their isolated forms. For the former, we use a normalized helical propensity (see Methods), HPNA, defined as HPa.a./NAa.a. where HPa.a. is the helical propensity of a given amino acid type and NAa.a is natural abundance of that amino acid (**Table 1**). Both HP_a.a._ and NA_a.a._ are probabilities, where HP_a.a_ is defined as the fraction of a certain amino acid type in TP-DB and NA_a.a._ is the fraction of the same type of amino acid (https://web.expasy.org/protscale/pscale/A.A.Swiss-Prot.html) from the based on the ExPASy tool “ProtScale”^36^. A log-odds ratio, log (HP_NA_) or log (HP_a.a._/NA_a.a._) is found for each amino acid type, and the reported helical propensity for each peptide in TP-DB is the sum of log (HPNA) for every constituent residue in this peptide (**Table 2**). The second metric is the contact number, defined as the average number of amino acids, represented by corresponding Cα atoms, contacting a helix of interest within 15Å from it. The lower contact number of a helix in its parent protein suggests a higher probability for it to stay in the helical form in isolation.

### 3.2. Anti-FLAG antibody is repurposed for pathogen diagnosis

To find specific patterns in a biological sequence while inferring homology, Pattern Hit Initiated (PHI)-BLAST has been the main (if not only) bioinformatics tool to perform such^18^. PHI-BLAST was designed to address evolutionary relevance of two proteins carrying the same or similar motifs, so it renders results considering both the pattern match and local sequence similarity with statistically significance. To showcase the prowess and disparate utility of the TP-DB, we demonstrate below how a purification tag (herein FLAG-tag) and its antibody (herein M2 antibody) can be repurposed into a diagnostic kit for detecting human pathogens. The relevant experimental methods are listed in Supplementary Materials. FLAG-tag is known to have a sequence DYKDDDDK^37^ with the pattern D-Y-K-x-x-[DE] being experimentally confirmed as the main affinity determinant motif^38^. “DYK” here is the strongest determinant while the last D (or E) is of the secondary importance. We would like to search TP-DB for proteins that (i) are not included in the PHI-BLAST search results and (ii) belong to proteins in human pathogens but not in non-pathogenic bacteria.

#### 3.2.1. Confirmed by ELISA and Western Blot, HP-1 is Recognized by anti-FLAG M2 Antibody Specifically at a stretch containing the D-Y-K-x-x-[DE] motif

When querying the pattern “D 0 Y 0 K 2 D/E” against the TP-DB, we can find totally 93 sequences with unique 19 sequences. While using PHI-BLAST to search NCBI’s non-redundant protein sequences (nr) with the pattern D-Y-K-x(2)-[DE] (in PROSITE format) and FLAG sequence “DYKDDDDK”, we could not find any sequence with an E-value less than 100. On the other hand, among the 19 TP-DB-identified sequences, HP-NAP, containing the sequence “**DYK**YL**E**”, can be found in *Helicobacter pylori* strain 26695 (*accession no.* AE000543, ATCC) but not in regular non-pathogenic bacteria. If the anti-FLAG M2 antibody can recognize this randomly selected pathogenic protein for its containing a FLAG-determinant motif (herein D-Y-K-x-x-[DE]), we can potentially use the same concept to repurpose the original use (say, purification) of any known pair of antibody and antigen recognition motif (say, M2 antibody and D-Y-K-x-x-[DE]) into a new use, herein the potential diagnostic kit for human pathogens. To our heartfelt delight, the anti-FLAG M2 antibody was indeed found to be capable of recognizing HP-NAP as analyzed by ELISA (**Fig. 2A**) and Western blot (**Fig. 2B**). Further analysis showed that M2 antibody detected maltose-binding protein (MBP)-tagged HP-NAP_R77-E116_ but not MBP (**Fig. 2B**), indicating that M2 antibody binds to the polypeptide fragment of HP-NAP containing residues Arg77 to Glu116, which contains the antigen recognition motif, **DYK**YL**E,** as shown in **Fig. 2C**.

**Fig. 2.**
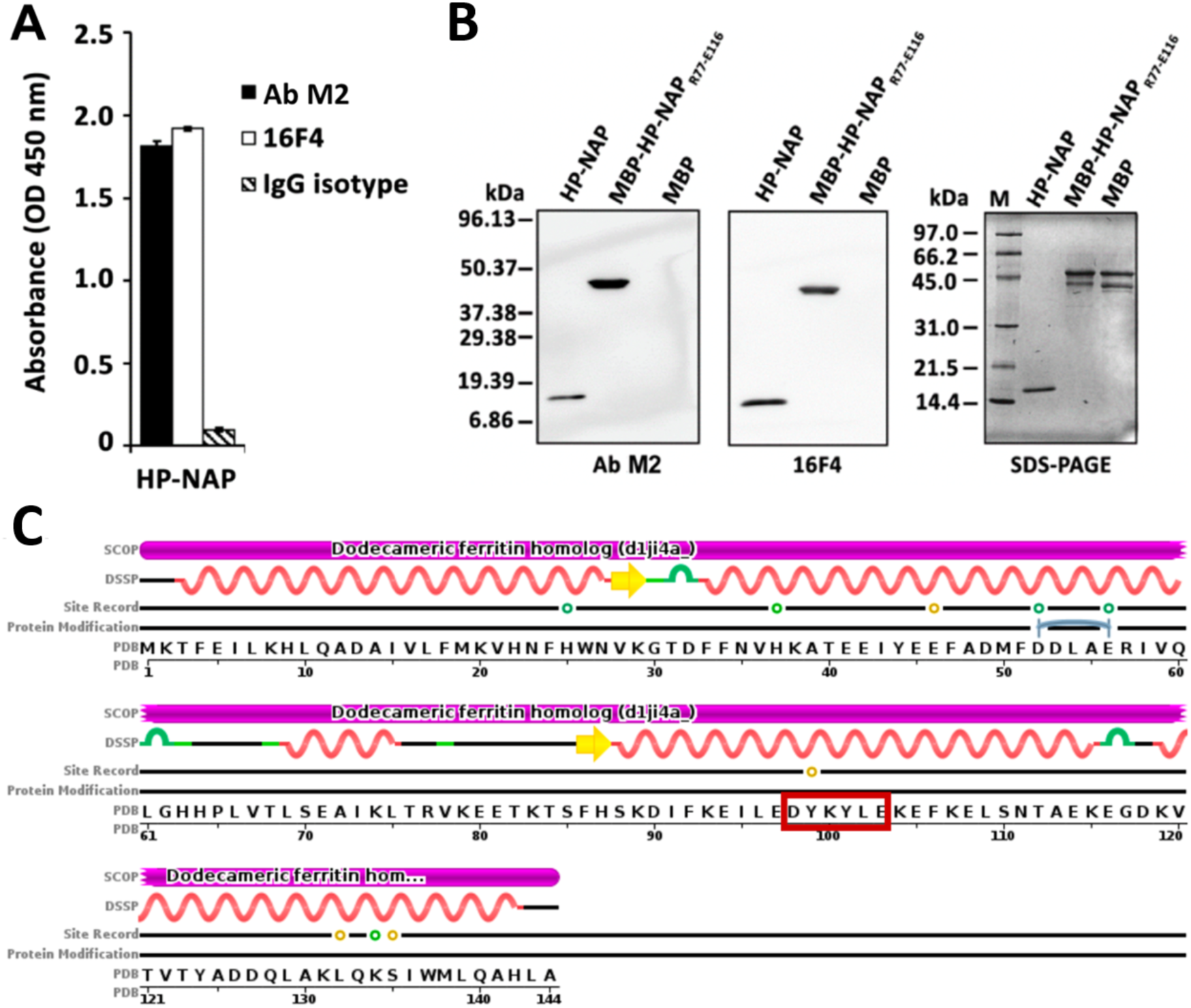
Detection of HP-NAP by anti-FLAG M2 antibody. (**A**) Detection of HP-NAP by anti-FLAG M2 antibody using recombinant HP-NAP-based ELISA. Recombinant HP-NAP was purified by a small-scale DEAE Sephadex negative mode batch chromatography. Purified HP-NAP as an antigen was coated on an ELISA plate as described in Methods. The anti-FLAG M2 antibody, its corresponding mouse IgG antibody, and the antibody 16F4 against HP-NAP, as positive control, were subjected to recombinant HP-NAP-based ELISA. The result is expressed as absorbance at OD_450 nm_. Data were presented as the mean ± S.D. of one experiment in duplicate. Similar results were obtained in three independent experiments. (**B**) Detection of recombinant HP-NAP by anti-FLAG M2 antibody using Western blot analysis. Recombinant HP-NAP was purified by two consecutive gel-filtration chromatography. MBP-tagged HP-NAP_R77-E116_ and MBP were partially purified by the MBPTrap HP column using ÄKTA Purifier. These purified proteins, 1 µg each, were subjected to SDS-PAGE on a 15 % gel and Western blot analysis with anti-FLAG M2 antibody and the antibody 16F4. Molecular masses (M) in kDa are indicated in the left of the blots and the gel. Similar results were obtained in two independent experiments. **(C)** The sequence of HP-NAP. Note that “DYKYLE”, highlighted in the red box, matches the pattern D-Y-K-x-x-[DE] and is included in the R77-E116 stretch. The image is download from PDB (code:1JI4) with minor modifications.

As a result, the antibody (herein M2 antibody) of a purification kit can be *repurposed into a diagnostic kit* for detecting human pathogens, herein *H. pylori*, through recognizing native HP-NAP by ELISA or HP-NAP monomer as a 17 kDa band by Western blot.

### 3.3. α-Helical Antimicrobial Peptide (AMP) Design

Pan-drug-resistant bacteria is no less a threat to mankind than viral infections that could cause a pandemic^39^. A discouraging forecast has suggested that by 2050, 10 million people could die from these super bugs each year, with half of them in Asia^40^. Antimicrobial peptides (AMPs) that target bacterial lipid membranes, evolving at a much slower pace than bacterial protein receptors, appear to be a promising alternative for antibiotics. There have been substantial efforts made to study AMPs’ insertion mechanism^26, 27, 41, 42^ and analyze the determining characteristics for their efficacy^43, 44^. To join these collective efforts and showcase the good use of TP-DB, we first examined in details the insertion processes of two previously reported helical antimicrobial peptides (AMPs), W3_p1 (sequence: KK **W**RK **W**LK **W**LA KK^45^) and W3_p2 (sequence: KK **W**LK **W**LK **W**LK KK^2^) by MD simulations. The two amphiphilic AMPs have 3 equally spaced tryptophans at their hydrophobic face, while positively charged lysines/arginines are on the other side, which is a typical feature and requirement for helical AMPs. The goal of the simulations is to understand each residue’s physicochemical role in the bacteria-membrane insertion process while earlier reports^2,45,46,47,48^ have suggested the deeper a peptide penetrates the lipid membrane, the higher bactericidal potency it can have.

With microsecond simulations, we revealed types of contact modes and time-resolved atomic interaction between AMPs and lipid bilayers before AMPs’ final insertion into the bacterial membrane (see **Movie S1 and Fig S2**). Taking W3_p1 for example, as shown in Figure 3, lysine and arginine with long positively charged side chains preferentially interact with the phosphate groups in PC and PG (see **Fig S1**). These interactions not only “raise” some interacting lipid molecules out of the upper leaflet but also push away these phosphate groups due to the long arms of these positively charged side chains. This creates what we call “hydrophobic cradle” which is a temporary vacuum created due to the aforementioned Lys/Arg-lipid interaction where at the bedding of the cradle lie the exposed aliphatic lipid tails whose head groups have been pushed either side way or upward. One can see such cradles form at 46, 673, 678 and 715 ns in **Figure 3**. This provides an important opportunity for adjacent tryptophan residue to flip in to enjoy such a hydrophobic interaction only when the orientation is right. Either W3 or W9 (see 46 ns in **Fig 3**) close to N- or C-terminal double lysine residues can be the first tryptophan to take advantage of these hydrophobic cradles for insertion. However, W9 insertion near the C-terminus is transient (starting right in the beginning, at 10 ns, drifting away at 46 ns, resuming the insertion at 93 ns but then drifting away again soon after) (**Movie S1 and** **Fig 3**). It cannot be maintained because the horizontal insertion pops the other two tryptophan up or side way toward the bulk solvent. On the other hand, W3-initiated insertion benefits not only from the transiently formed hydrophobic cradles but also from the downward facing N-terminus (see 530 ns, 715 ns and thereafter in Fig 3) that further brings W6 to contact the deeper part of a lipid (e.g. 715 ns and thereafter). The N-terminus-rather than C-terminus-initiated penetration is consistent with aforementioned experimental results^49^ and also common to most of the membrane anchoring signals^50^. N-terminus seems to better interact with the phosphate groups of lipids (e.g. at 678ns) and transiently interact with the lipid tails better than the C-terminus.

**Fig. 3.**
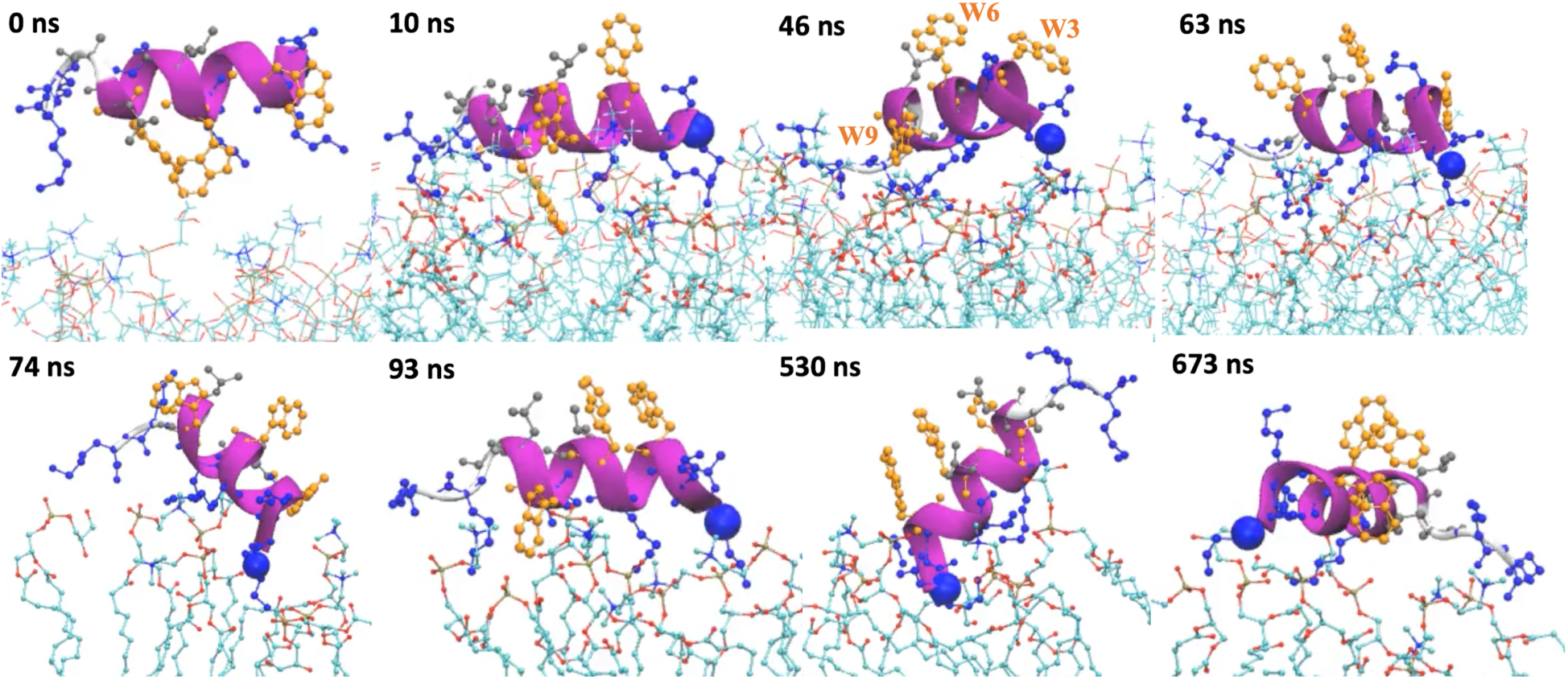
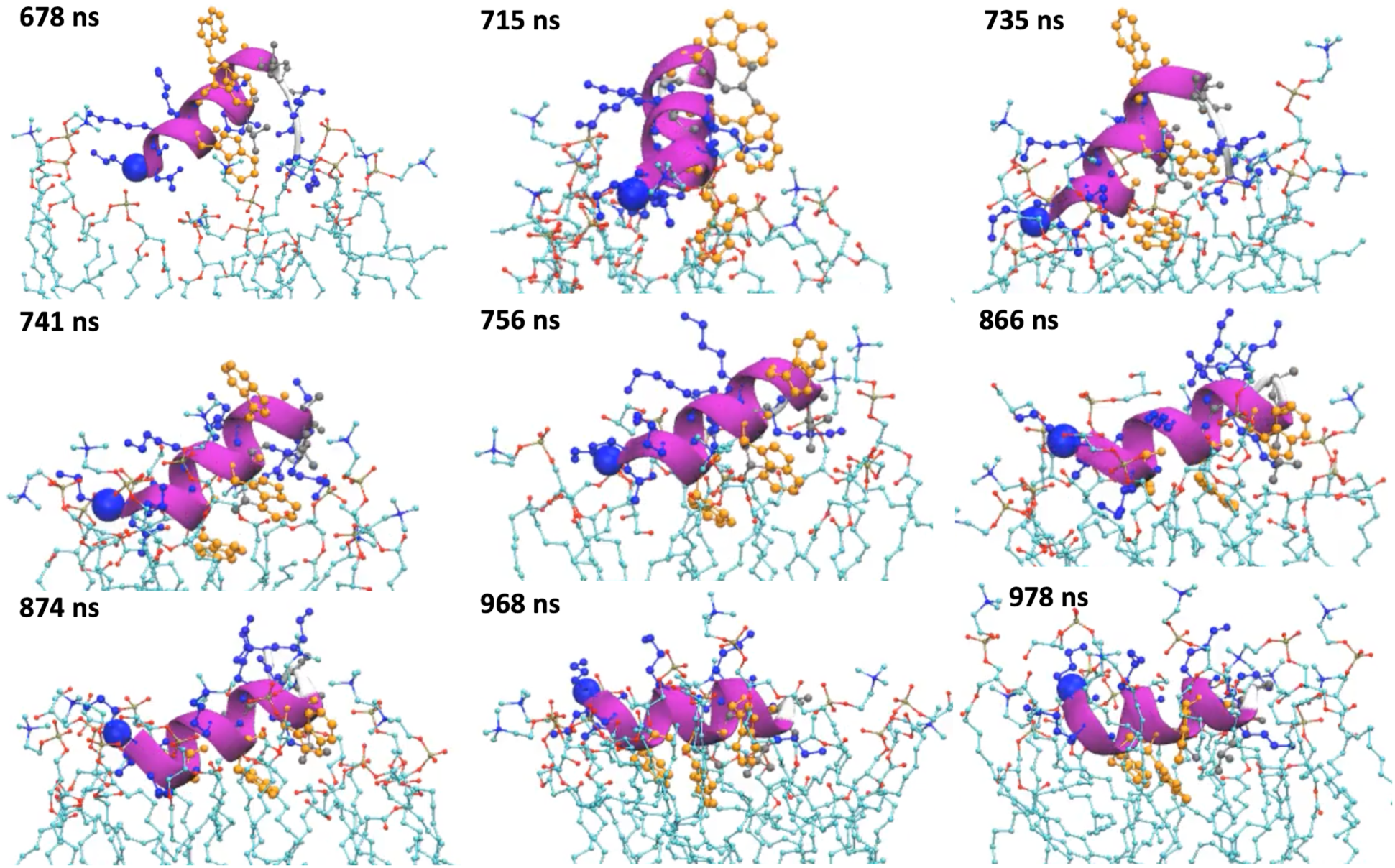
Mechanistic details of W3_p1’s insertion into the bacterial membrane revealed by 1 µs MD simulations. The Cα atom of the first residues (N-terminus) is highlighted with a blue VDW ball, from which the three tryptophan residues, shown in golden ball-and-stick, are W3, W6 and W9 (see also the panel of 46 ns). The positively charged residues in the AMP are represented in blue CPK. Starting from the 74th nanosecond, only the lipids staying within 3 Å from the AMPs are shown while part of their aliphatic tails can be removed for clarity. The heavy atoms of the lipids are represented in thin CPK; O, P and C atoms are colored in red, brown and cyan, respectively. The N atoms in the choline groups of POPC lipids that engage cation-π interactions with the tryptophan are colored in blue.

As W3 starts to insert, its irreplaceable chemical importance in membrane insertion becomes immediately apparent in these time-resolved snapshots. As shown at 678 ns in Figure 3, it can first interact with the choline groups of PC lipids through the cation–π interaction. Then, it can interact with the phosphate groups (715 ns) or deeper side of the glycerol moiety (735 ns) in the same or different lipids via hydrogen bonds. Eventually W3 reaches and interacts stably with lipid tails due to the hydrophobic interaction (741 ns onward). As W3 reaches the lipid phosphate groups or glycerol moieties, W6 can start to interact with the choline group of the same lipid (e.g. at 715 ns). When W6 starts to reach the phosphate group (735 ns) and lipid tails (741 ns), the originally vertical W3_p1 starts to lean down with the N-terminus still diving a bit deeper than the C-terminus, which brings W9 closer to the lipid on the side way. At 756 ns, we see that W9 starts to interact with two choline groups of PC lipids. As W9 starts to sink deeper to interact with the phosphate group of a lipid (e.g. at 866 ns), to allow W3 and W6 being still stably anchored in the lipid tails, the N-terminus starts to open up and becomes less helical (866 ns onward), which helps split W3 and W6 further apart and bring W9 to rotate further downward and better align W6 and W9 on the same side. This also synchronizes with positively charged residues being upward facing and sticking their side chains toward the bulk solvent (968 ns onward).

In summary, W3_p first interacts with lipids via electrostatic interaction with the phosphate groups of the lipids popping up tryptophans in the bulk solvent. As N-terminus dives downward in the hydrophobic cradle, created by the long positively charged side chains to push the lipid heads away and expose their lipid tails, the W3 is brought together to interact with the deeper part of the lipids, which makes the AMP stand vertically on the lipid surface. As W6 further crawls in and interacts with phosphate groups of the lipids, or deeper, the AMP starts to lean down with N-terminus in turn to become “loopy” and less helical, which helps W9 be better aligned with W6 and insert deeper. As also evident in the Movie S1, within the 1 microsecond, all the 3 tryptophan residues in W3_p1 are inserted into the bacterial membrane while W3_p2, which is strictly helical, has only two tryptophan residues (W3 and W6) inserted into the membrane (especially after 800 ns). In short, these AMPs insert the bacteria membrane heavily relying on the *three tryptophan anchors* equally spaced with two amino acids.

Knowing the aforementioned insertion process and the importance of the pattern **“**W**W**W” (or [W 2 W 2 W] as the input query in TP-DB) in a helical structure, we are interested in finding other helical peptides in TP-DB with the following three characteristics - (1) containing the W2W2W motif, (2) maintaining helicity in isolation (with relative high HP) and (3) enriched with positively charged residues, so that the corresponding part in W3_p1 or W3_p2 can be replaced by the TP-found helical stretches (**Table 2**) to preserve the terminal lysine anchors.

105 helical peptides from different PDB IDs (and chain IDs) containing 11 unique **“**W**W**W**”** motifs can be found through the query [W 2 W 2 W] (**Table 2**). The peptide **“**WKCWARWRL” (the 5th in the #U column) having the most positively charged residues and a relatively high HP is chosen as our top AMP candidate. The peptide is derived from the mycobacterium tuberculosis Zinc-dependent metalloprotease-1 (Zmp1) (PDB: 3ZUK), which is a soluble protein and does not associate with the membrane. The peptide sits at the relatively outerior side of the protein but is not wide open to the bulk solvent. The flanking double Lys residues are further added to extend this motif into a full-fledged AMP candidate. This newly designed AMP, named as W3_db5, is then investigated by MD simulations and synthesized for the subsequent bactericidal experiments.

Contrasting to the simulation results of W3_p1 and W3_p2, W3_db5 has only one Trp (W3) inserted into the bacterial membrane by the end of 1 microsecond simulation (Movie S1 and Fig S2). To further examine the bactericidal and antifungal activities of these AMPs, we measure the minimal inhibitory concentration (MIC_90_) at which the growth of 90% of the pathogenic fungus *C*. *albicans* SC5314 and *E. coli* ATCC 25922 can be inhibited. Experiments are performed in pentaplicate and MIC_90_ is determined as the majority results of the 5 replicates (see Methods). It is worth mentioning that we could reproduce the reported MIC_90_ values for three published AMPs when using the same culture protocol to culture *E. coli* ATCC 25922. They are CM15, Anoplin-R5KT8W and FK13 with reported MIC values of 4^51^, 15.13^52^ and 27.5^53^ µg/ml, respectively, while we obtained 6.25, 12.5 and 25 µg/ml for the same. Our results (**Table 3**) show that the newly designed AMP W3_db5 (KK **W**KC **W**AR **W**RL KK) has an anti-fungal MIC_90_ of 3.75 µg/ml, which is twice better than that of W3_p1 (7.5 µg/ml) and W3_p2 (7.5 µg/ml), while it is a level lower in its bactericidality with a MIC_90_ of 30 µg/ml than that of W3_p1 (15 µg/ml) and W3_p2 (15 µg/ml). The latter result is consistent with the MD-based natural insertion results (Fig S2).

**Table 3.**
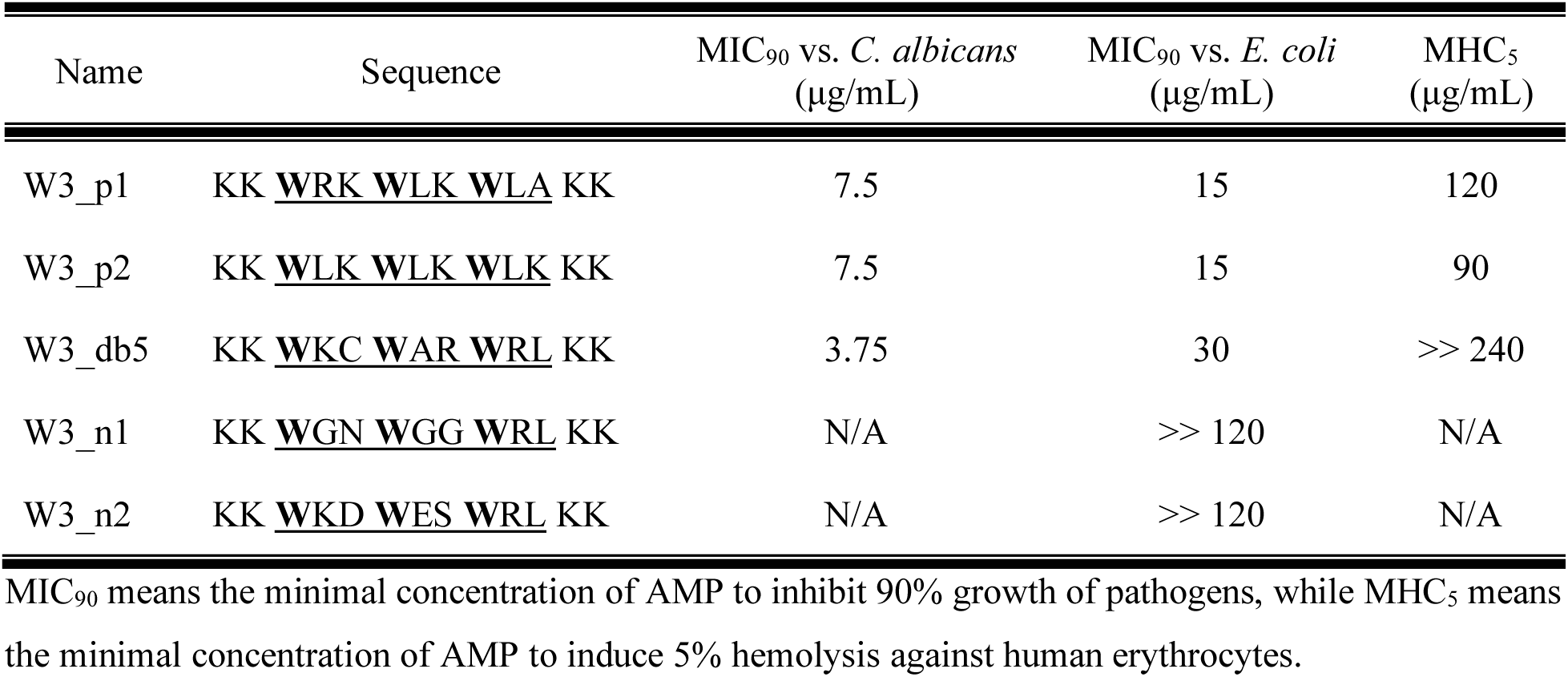
The anti-fungal, antibacterial activity and hemolytic toxicity of TP-DB-designed AMP (W3_db5) and other AMPs containing the “W2W2W” motif.

| Name | Sequence | MIC<sub>90</sub> vs. *C. albicans*<br>(µg/mL) | MIC<sub>90</sub> vs. *E. coli*<br>(µg/mL) | MHC<sub>5</sub><br>(µg/mL) |
| --- | --- | --- | --- | --- |
| W3_p1 | KK <u>WRK WLK WLA</u> KK | 7.5 | 15 | 120 |
| W3_p2 | KK <u>WLK WLK WLK</u> KK | 7.5 | 15 | 90 |
| W3_db5 | KK <u>WKC WAR WRL</u> KK | 3.75 | 30 | >> 240 |
| W3_n1 | KK <u>WGN WGG WRL</u> KK | N/A | >> 120 | N/A |
| W3_n2 | KK <u>WKD WES WRL</u> KK | N/A | >> 120 | N/A |

MIC<sub>90</sub> means the minimal concentration of AMP to inhibit 90% growth of pathogens, while MHC<sub>5</sub> means the minimal concentration of AMP to induce 5% hemolysis against human erythrocytes.

Although our newly designed W3_db5 has similar anti-bacterial and anti-fungal activities as its mother templates’, it has a surprisingly low cytotoxicity as compared with the template AMPs. As shown in Figure 4, at the concentration of 240 µg/ml, W3_p2, W3_p1 and W3_db5 lyse 20.3%, 13.0% and 0.6%, respectively, the red blood cells (RBCs). W3_db5 is at least 20-fold less hemolytic than W3_p1 and W3_p2 (at least 12-fold less hemolytic at 120 µg/ml); in other words, we can use 20-fold higher concentration than W3_db5’s MIC to still have a smaller cytotoxicity than its W3 templates. In Table 3, we also list AMPs’ minimal hemolytic concentration (MHC5) to lyse 5% of the RBCs, where W3_db5’s MHC5 can be much larger than 240 µg/ml, which implies even a high concentration of W3_db5 can still be potentially used in medication or personal hygiene products.

**Figure 4.**
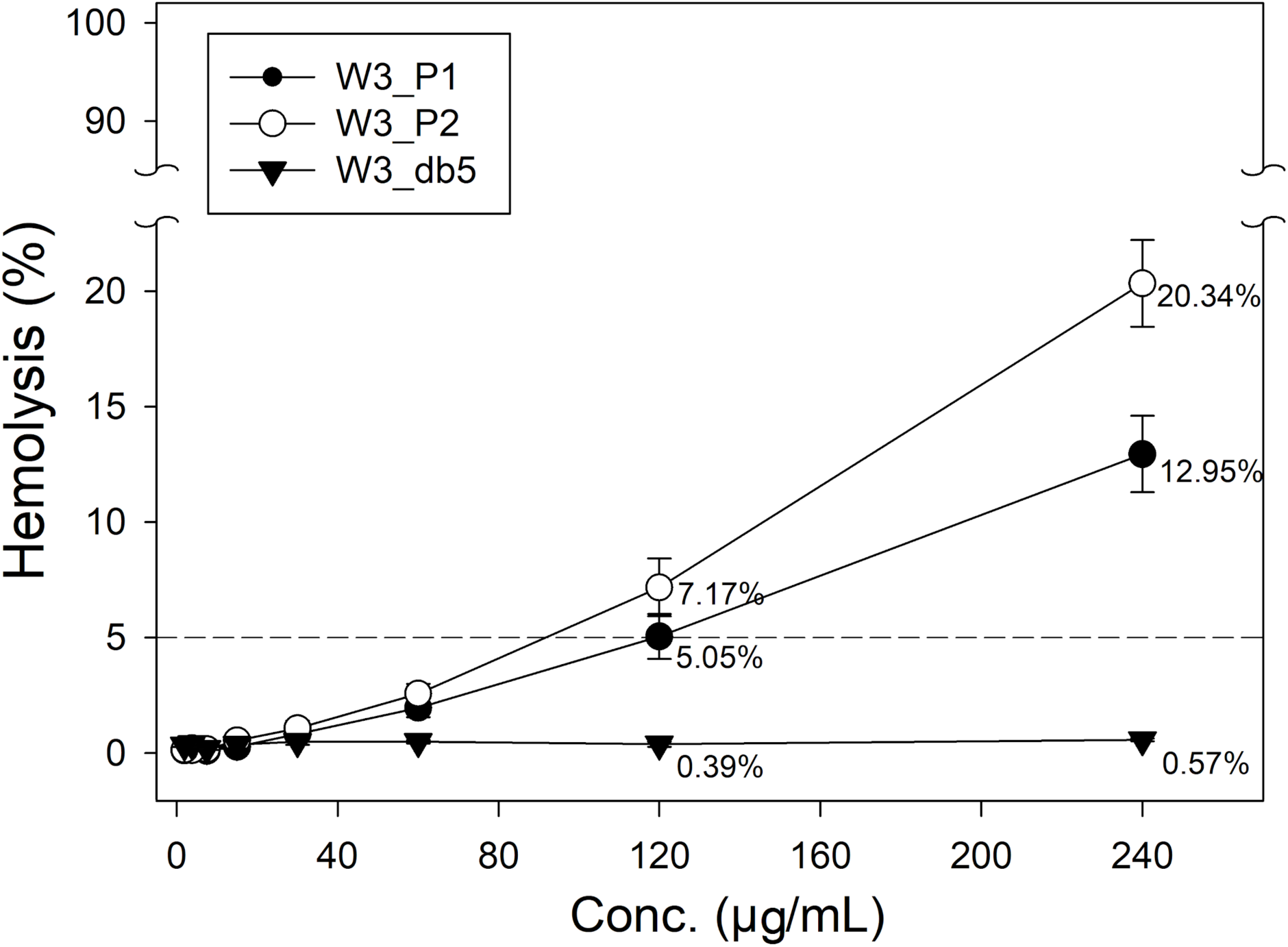
The hemolytic percentage as a function of AMP concentrations for W3_p1, W3_p2 and W3_db5. W3_db5 has a much lower hemolysis of red blood cells (RBCs) than that of W3_p1 and W3_p2. The dash line indicates the 5% hemolytic threshold that defines MHC5. Data are presented by the mean ± S.D. from three repeats.

To further validate the use of TP-DB, as a control study, we borrow the conventional PHI-BLAST to find a sequence homologous to W3_p1 and containing the motif “W2W2W”. Here we have to note that PHI-BLAST has to take two input to initiate a search - a protein sequence S and a regular expression pattern P occurring in S. Given the two inputs, PHI-BLAST helps finding protein sequences both containing an occurrence of P and are homologous to S in the vicinity of where the pattern P occurs. Here, P as “W-x(2)-W-x(2)-W” and S as the sequence of W3_p1 are used to search the PDB via the NCBI PHI-BLAST website. A single chain protein (PDB: 4F54) from the DUF4136 family with a relatively high resolution (1.60 Å) contains a stretch of “**W**GN **W**GG **W**”, located at the end of a helix connecting toward a beta-sheet through a short loop. We synthesize a peptide “KK WGN WGG WRL KK”, coined as “W3_n1”, which replaces the “**W**KC **W**AR **W**” part in W3_db5 with the “**W**GN **W**GG **W**”. When examining its MIC, we fail to find its MIC_90_ given the concentration range we have tested (the highest at 120 µg/ml) (see **Table 3**), suggesting not any non-helical “W2W2W” sequence can work as a functional AMP motif. As a second control study, we take another helical “W2W2W” stretch in **Table 2**, herein the 8th unique peptide “**W**KD **W**ES **W**”, to a new “AMP”. That means we replace the “**W**KC **W**AR **W**” part in W3_db5 with “**W**KD **W**ES **W**” to create the peptide “KK **W**KD **W**ES **W**RL KK”, which is termed as “W3_n2”. The “**W**KD **W**ES **W**RL” stretch has one less positively charged residue and two more negatively charged residues (D and E) as compared with the “**W**KC **W**AR **W**RL” motif in W3_db5. Again, we do not find its MIC_90_ within the concentration range we have tested (the highest at 120 µg/ml) (see **Table 3**), which suggest a helical stretch containing the W2W2W motif does not necessarily guarantee its bactericidal activity if there are not enough positive charges in the designed AMPs. Note that the TP-DB-found W3_db5 has the same number of positive charges as its W3 templates (i.e. W3_p and W3_p2), while W3_n2 could be 3 positive charges short to become an AMP. In the MIC_90_ experiments, W3_n1 and W3_n2 respectively kill 2.5% and 9.5% of the *E. coli* ATCC 25922 at a concentration of 30 µg/ml, which is the MIC_90_ value of W3_db5 for the same bacteria.

It should be noted that the original protein (mycobacterium tuberculosis Zinc-dependent metalloprotease-1; PDB ID: 3ZUK) that contains this potent stretch “**W**KC **W**AR **W**RL” in W3_db5 is not a transmembrane or membrane-bound protein; the helical stretch is situated close but not fully exposed to the water-protein interface. Therefore, TP-DB brings an interesting opportunity for researchers to put together structurally resolved elements, matching a desired pattern, for the rational design of new therapeutics, herein showcased by a new class of AMP that is 6 point mutations away from its original templates - W3_p1 and W3_p2.

From the design point of view, these 6 point mutations do not seem to change much the required insertion thermodynamics^54^ in the new AMP, W3_db5, which can still penetrate the peptidoglycans, reach the lipid-water interface, stay amphiphilic and helical, and eventually insert into the cell membrane. This is difficult to achieve by mutagenesis experiments or MD simulations without the guidance of TP-DB.

### 3.4. Design Peptide Inhibitors for Tumorigenic PPI

Our statistics in **Table 1** also reveal the interesting fact that alanine is indeed a safe choice for point mutation while still maintaining the integrity of a helix (see HPNA). However, with the pattern-based search engine (see Methods section 2.2) made available in TP-DB, one can do point mutation(s), by choosing the residues suggested by TP-DB, in a helix to abolish or enhance (see Supplementary Material) its interaction with certain drugs, peptides or proteins while still maintaining the helicity of the helix, as well as to abolish the catalytic function of a residue on an *α*-helix while maintaining its binding affinity with certain substrate or ligand in the scenario where a structural biologist co-crystalizes an enzyme with its cognate substrate. In the Supplementary Material, we demonstrate an *in-silico* study on how peptide blockers can be designed by TP-DB to block a helix-helix interface in order to suppress the late-stage hepatocellular carcinoma.

## 4. Conclusions

The pattern-based search engine without relying on sequence homology, realized in the newly introduced TP-DB, has been shown to provide a new avenue to the re-use or re-assembly of structured peptide fragments into new therapeutics or diagnostic kits. New therapeutics can therefore be designed not only with known functional sequence motifs but also with desired secondary structures. We also foresee its future extension to incorporate search engines for beta-sheet elements containing specific sequences and/or patterns to meet desired physicochemical traits with a design purpose.

## Supporting information

Supplementary Results

## Acknowledgements

E.O.S. acknowledges financial support from MOST and Taiwan International Graduate Program, Academia Sinica, Taipei, Taiwan. We thank vast computational resources for this project provided by High Performance Computing Infrastructure (HPCI), Japan and National Center for High-performance Computing (NCHC) of National Applied Research Laboratories (NARLabs) of Taiwan. This work was supported by the Ministry of Science and Technology, Taiwan (103-2627-M-007-001 and 104-2113-M-007-019 to L.W.Y; 104-2311-B-007-003 and 108-2311-B-007-004 to H.W.F) and the research program of National Tsing Hua University (104N2052E1 to H.W.F).

## Supplementary Results

### Chemical structures of POPC and POPG lipids

**Figure. S1.**
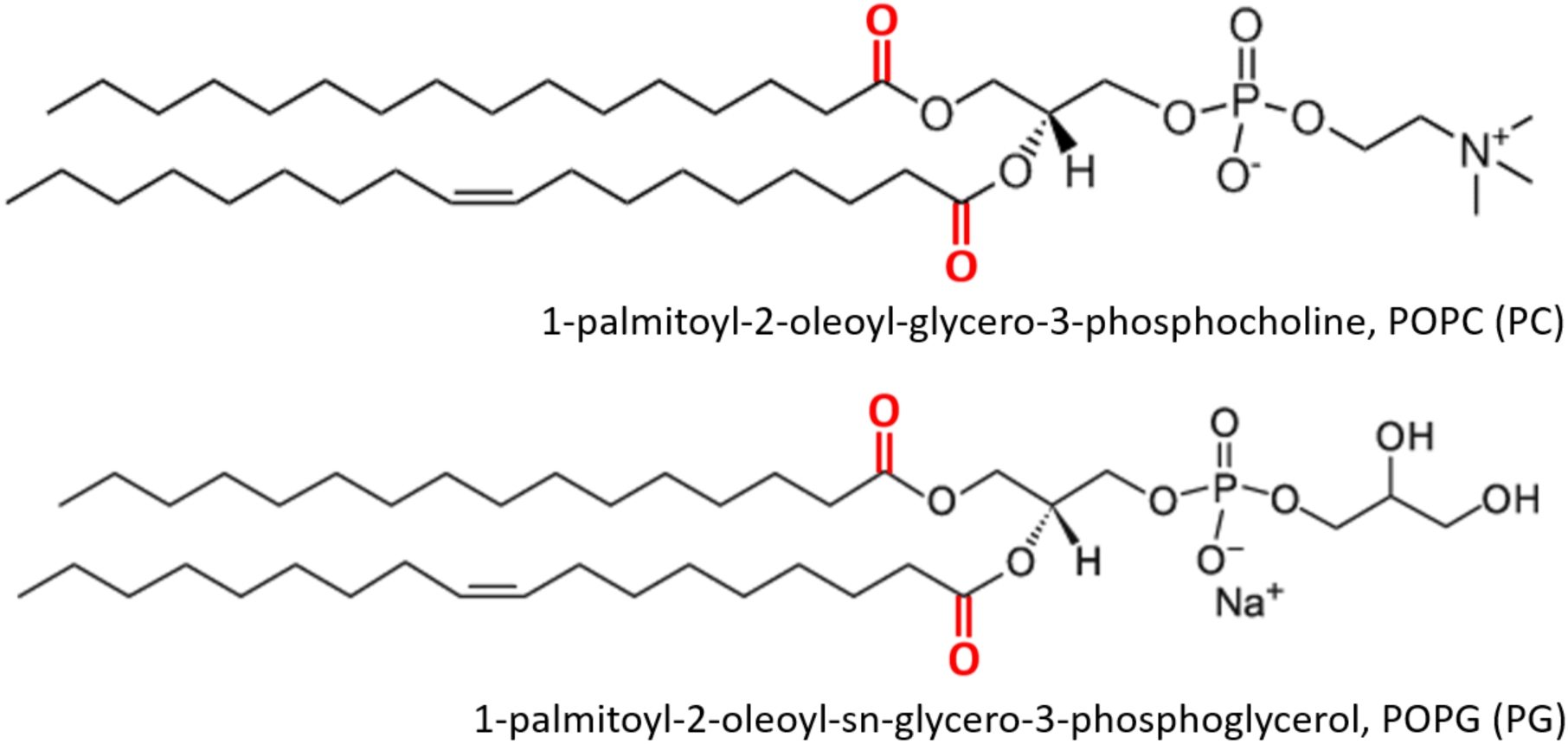
Chemical structure of POPC (PC) and POPG (PG). All images modified from Avanti® product page (Product ID: 840457 and 850457). The carbonyl oxygen in the lipids are highlighted in red, and these oxygen atoms are represented as red points in Fig. 3A. The atoms of the inserted AMPs below these oxygen atoms are considered to interact with the aliphatic tails of the lipids.

### Natural insertion of three AMPs into bacteria membrane

**Figure S2.**
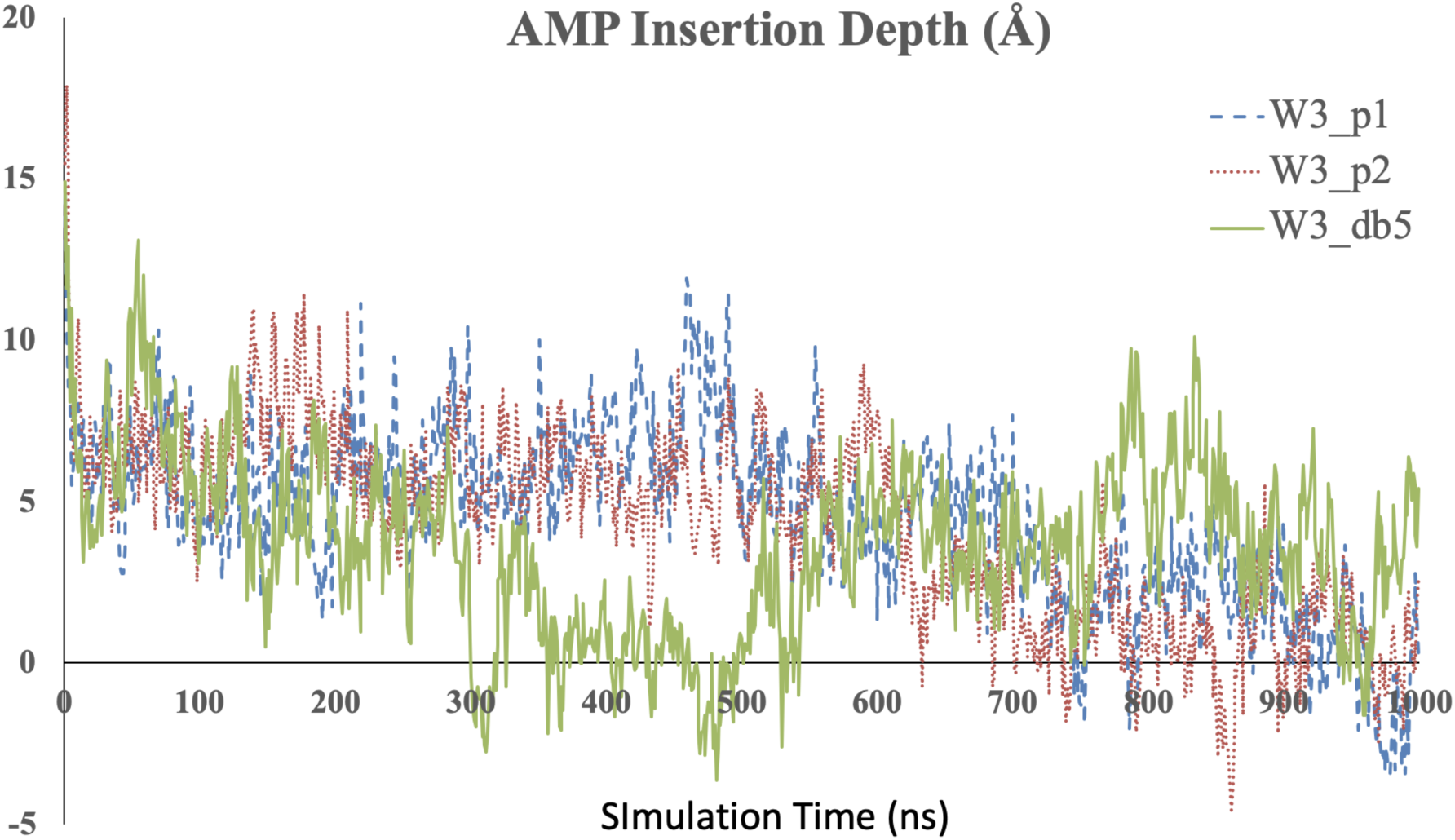
The relative position of the center of mass (COM) of AMPs to the COM of phosphor atoms in the upper leaflet consisting of 40 POPC/POPG (3:1) lipid molecules, mimicking bacteria membrane, is plotted against the simulation time. All the MD simulations were conducted for 1 microsecond (us) at body temperature and normal pressure (1 atm) using OpenMM^20^ with CHARMM36 forcefield^21, 22^.

## Supplementary Movie S1

All the three AMPs were first created as fully extended AMPs and they were then simulated in an NPT ensemble for 50 ns in the absence of lipids. The resulting snapshots of the AMPs were clustered into 5 groups based on their structural similarity using the “clustering” plug-in of the VMD software^31^. The representative conformation from the biggest cluster in each of the three AMPs was selected as the initial conformation to be simulated from time zero in an NPT ensemble in the presence of the already equilibrated PC/PG (3:1) membrane. At time zero, as seen in the video, all the peptides have been folded into a helical shape in the CHARMM36 forcefield^21, 22^. Within 1 microsecond, the tryptophan residues in W3_p1 had all three Trp and W3_p2 had two Trp (W3 and W6) inserted into the membrane (especially after 800 ns), while only one Trp (W3) in W3_db5 had been inserted into the membrane.

### Design of peptide inhibitor against Sgo1-PP2A interaction

Below we demonstrate how TP-DB can be used to design helical blockers to prevent disease-related protein-protein interaction.

In late-stage hepatocellular carcinoma (HCC), patients rely only on targeted therapy and chemotherapy, which were proved generally ineffective. Novel therapeutic targets for treating HCC are urgently needed. Researchers previously found that protein Shugoshin-1 (Sgo1), protecting the stability of centromeric cohesin at centromere and ensuring proper chromosome separation^55^, was up-regulated in HCC. Notably, hepatoma cells were found more sensitive to Sgo1 deficiency than normal hepatocytes^56^. These results could suggest that Sgo1 can be a potential therapeutic target for HCC. It has been known that phosphorylated-Sgo1 recruits protein phosphatase 2A (PP2A) to centromere during early mitosis^57^ and the Sgo1-PP2A complex sequestered SA2 subunit of cohesin from Plk1-mediated phosphorylation to maintain its stability^58^. Additionally, Sgo1-PP2A complex also maintains Sororin in a hypo-phosphorylated state and keeps the interaction of Sororin-Pds5 to counteract WAPL^57^. These regulations ensure centromeric cohesin tethering sister chromatids together, until the onset of anaphase (Fig S3). As a result, an inhibitor (possibly a peptide) that blocks the Sgo1-PP2A interaction could potentially suppress the rapidly growing hepatoma cells, susceptible to improper Sgo1-PP2A association, in mitosis.

**Figure S3.**
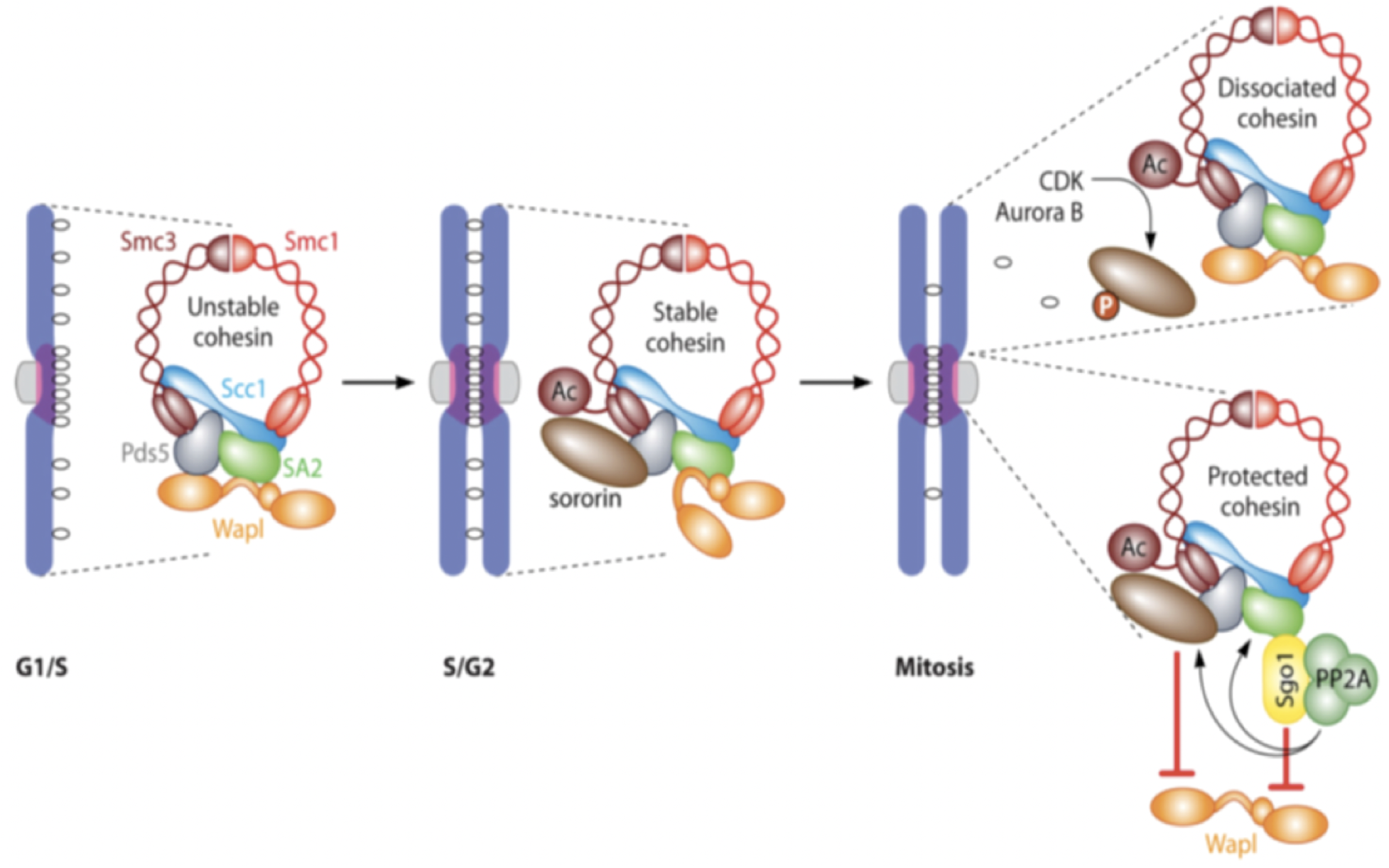
The overview of Sgo1-PP2A protecting centromeric cohesin. During mitosis, phosphorylation of SA2 and Sororin cause cohesin to dissociate from chromosome arm. Cdk1-mediated phospho-Sgo1 recruits PP2A at centromere to counteract phosphorylation of SA2 and Sororin and maintains centromeric cohesin to tether sister chromatids together. Image is adopted from Figure 3 in Marston’s review article^59^.

We noted that the complex of Sgo1-PP2A has been solved by x-ray crystallography (PDB ID: 3FGA) and the two molecules bind with each other through an helix-helix interface (one helix from each molecule)^11^. To design anticancer peptides as blocker against Sgo1-PP2A interaction, the helical stretch in PP2A (residues from 352-392 in chain B of PDB 3FGA), known to form the helix-helix interaction with another helix in Sgo1 (residues from 75 to 92 in chain D of PDB 3FGA) (Fig S4), is chosen as the candidate target for further investigation. Evidenced from the x-ray structure, the helix-helix interaction is visibly apparent to be the most important essential interacting elements at the protein-protein interface. As shown in Figure S3a, K374, G378 and Y381 in the helical stretch “KTIHGLIY” could be the anchoring residues on PP2A to mediate the Sgo1-PP2A interaction, which is supported by our calculation of energetics using *in silico* alanine scanning (Fig S4b).

**Figure S4.**
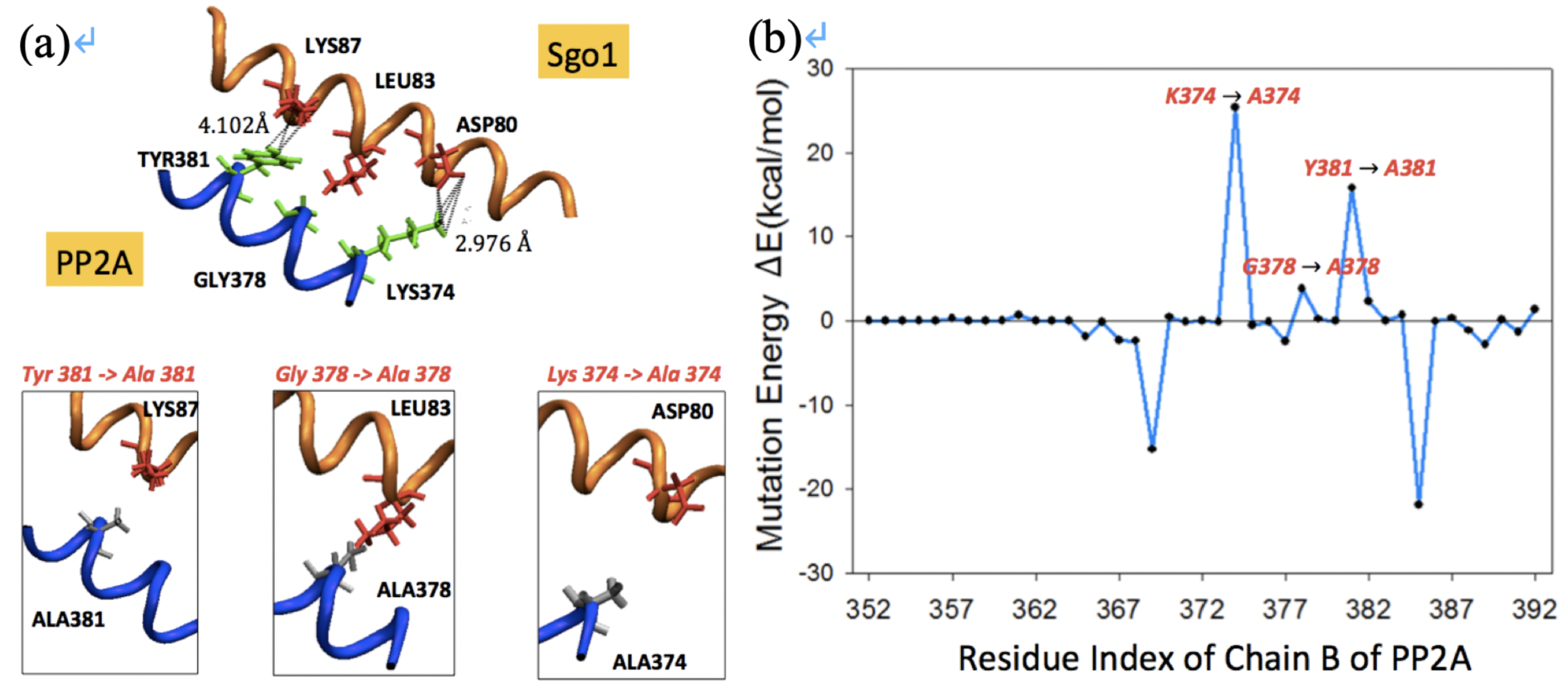
Protein-protein interaction (PPI) between Sgo1 and PP2A through helix-helix interactions. Panel **(a, upper)** shows only the PPI interface between Sgo1 and PP2A, while the rest of proteins (3FGA; Xu et al., 2009) are hidden for clarity. Sgo1-PP2A interaction is mediated by two helices, one from Sgo1 (PDB: 3FGA; V_75_KEAQDIILQLRKECYYL_92_ in chain D) and the other from PP2A (PDB: 3FGA; K_369_THMNKTIHGLIYNALK_385_ in chain B) . Computational alanine scanning for every residue in the PP2A helical fragment (from residue 352 to 392 of chain B) is carried out, and in panel (**b**) energetic contribution for each residue mutated into alanine is assessed (from residue 369 to 385 of chain B) where a large positive value (e.g. for K374A, G378A and Y381A) indicates the importance of the residues in binding, and a large negative value suggests mutation into alanine favors the binding. The non-bonded energy is evaluated by NAMD package (Phillips et al., 2005) using CHARMM 36 forcefield (Huang and MacKerell, 2013). Here in panel **(b)**, the mutation energy for the n-th residue, ΔEn, is defined as ΔEn = Yn - X0 where X0 is the non-bonded energy of original complex between Sgo1 and PP2A (for the 352-392 fragment), and Yn is the energy of the complex with the *n*-th residue in PP2A being mutated into alanine, after a short energy minimization.

To block the protein-protein interaction (PPI) between Sgo1 and PP2A, one could potentially modify part of the most essential interacting element in PP2A into a peptide that interacts with Sgo1 better than its template. As shown in Figure S4b, computational alanine scanning helps detecting a stretch from 370 to 384 in PP2A (chain B) dominantly contributing the affinity in the PPI of Sgo1-PP2A complex, especially from K374, G378 and Y381 (i.e. aforementioned essential interacting element in PP2A). We then used this helical stretch, or the extended one with its flanking region from residue 364 to 388, including two energetically favored point mutations, K369A and K385A, as the starting template.

To find the helical peptides that bind the interface helix of Sgo1 stronger than PP2A does, we search the TP-DB for PP2A-like helical peptides using the queries summarized in Table S1.

As detailed in Supporting Methods, we first used the pattern “**K***G**Y**” to search TP-DB and discover 69 unique PP2A-like helical peptides (Table S1 and S2); we then carried out molecular dynamics simulations (MD) to assess how these peptides interact with Sgo1 at the helix-helix interface. Helices H60, H54, H61, H33 and H29 (**Fig S5**) are energetically promising for further experimental confirmation. Additionally, we examined other sets of peptides that met the patterns consisting of three anchoring residues (“A***K***G” and “G**Y***A”) or consisting of at least four residues (“A***K***G**Y”, “K***G**Y***A”, “A***K***G**Y***A” and “A**/***K**/***G**/***Y**/***A”) in TP-DB. All the corresponding results obtained from the queries are illustrated in **Table S3** to **S8**. Among them only the “A***K***G**Y***A” pattern (according to Figure S3b) returns no result from TP-DB.

**Figure S5.**
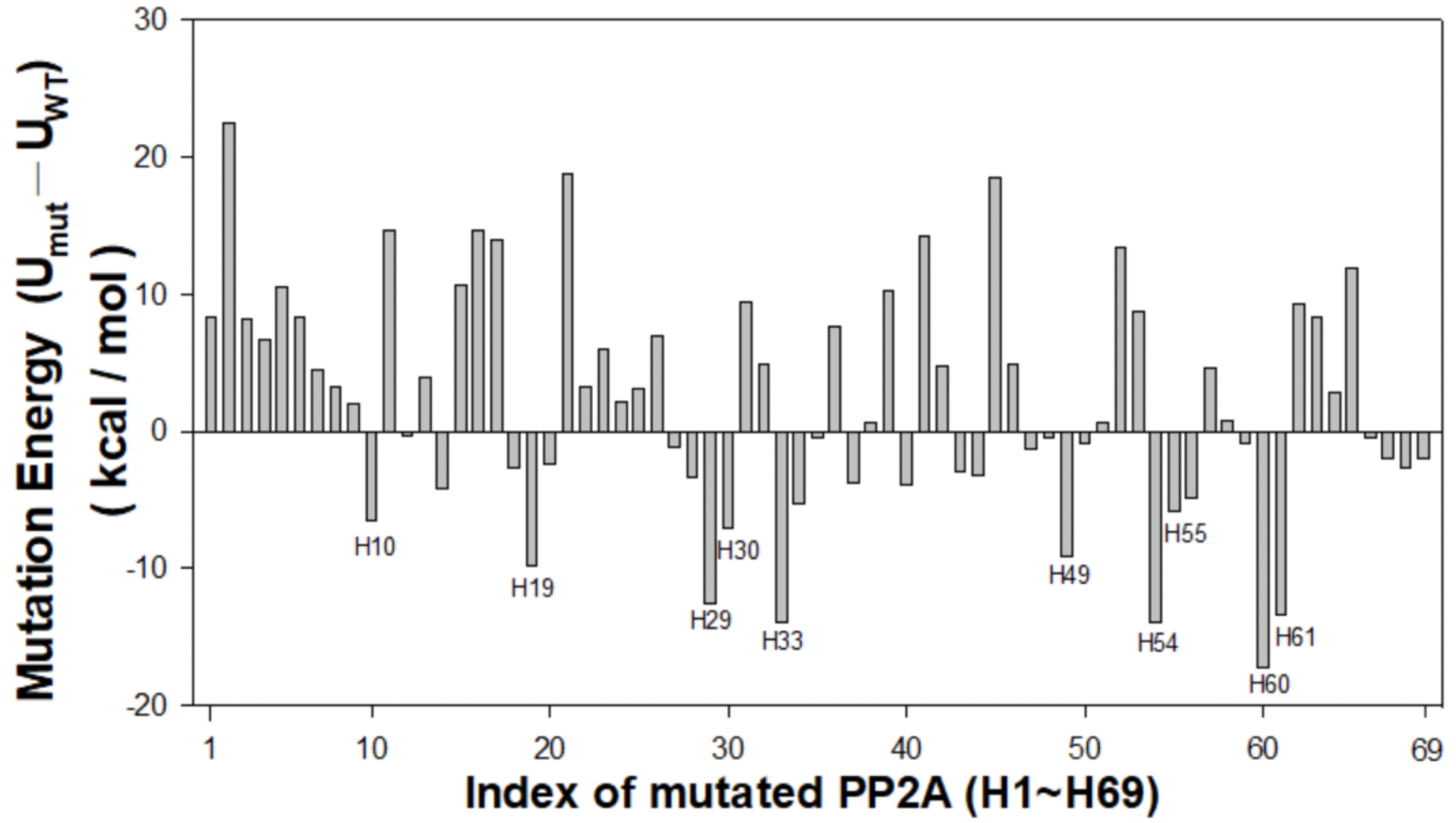
Interactions between Sgo1 helix and PP2A-like peptides. U_mut_ is the interaction energy between targeted Sgo1 helix and each of the 69 PP2A-like peptides (from H1 to H69) that match the pattern “K3G2Y”. In detail, the energy is calculated by replacing the residue T375, I376, H377, L379 and I380 in the “**K_374_**TIH**G**LI**Y_381_**” stretch of PP2A with the corresponding residues in each of the 69 pattern-matched (K3G2Y) helical peptides returned from TP-DB. UWT is the interaction energy between Sgo1 helix and PP2A. Larger difference of interaction energies (U_mut_ - U_WT_) suggest stronger binding of PP2A-like peptides with the Sgo1 helix than that between PP2A and the Sgo1 helix. Top ten values are further labeled in the graph.

**Table S1.**
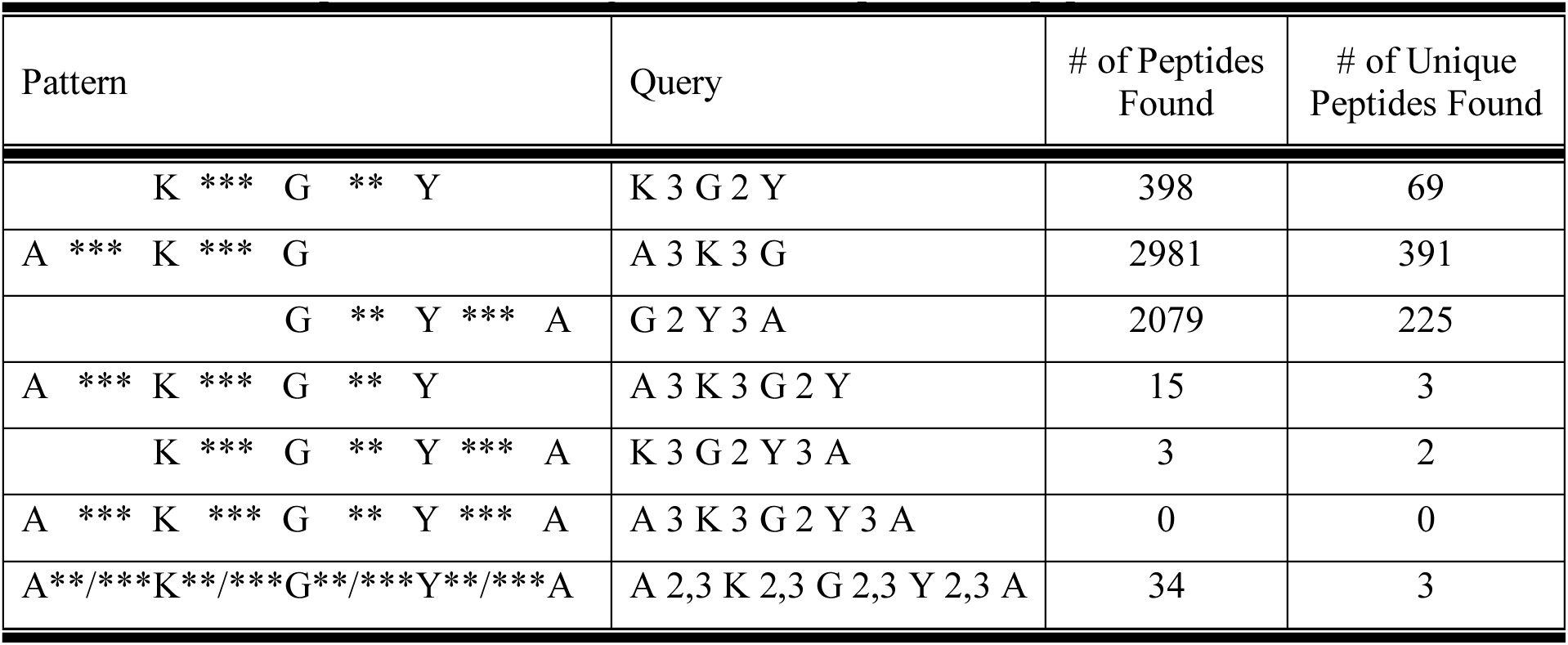
PP2A-like patterns searched against the developed helical peptide database

**Table S2.**
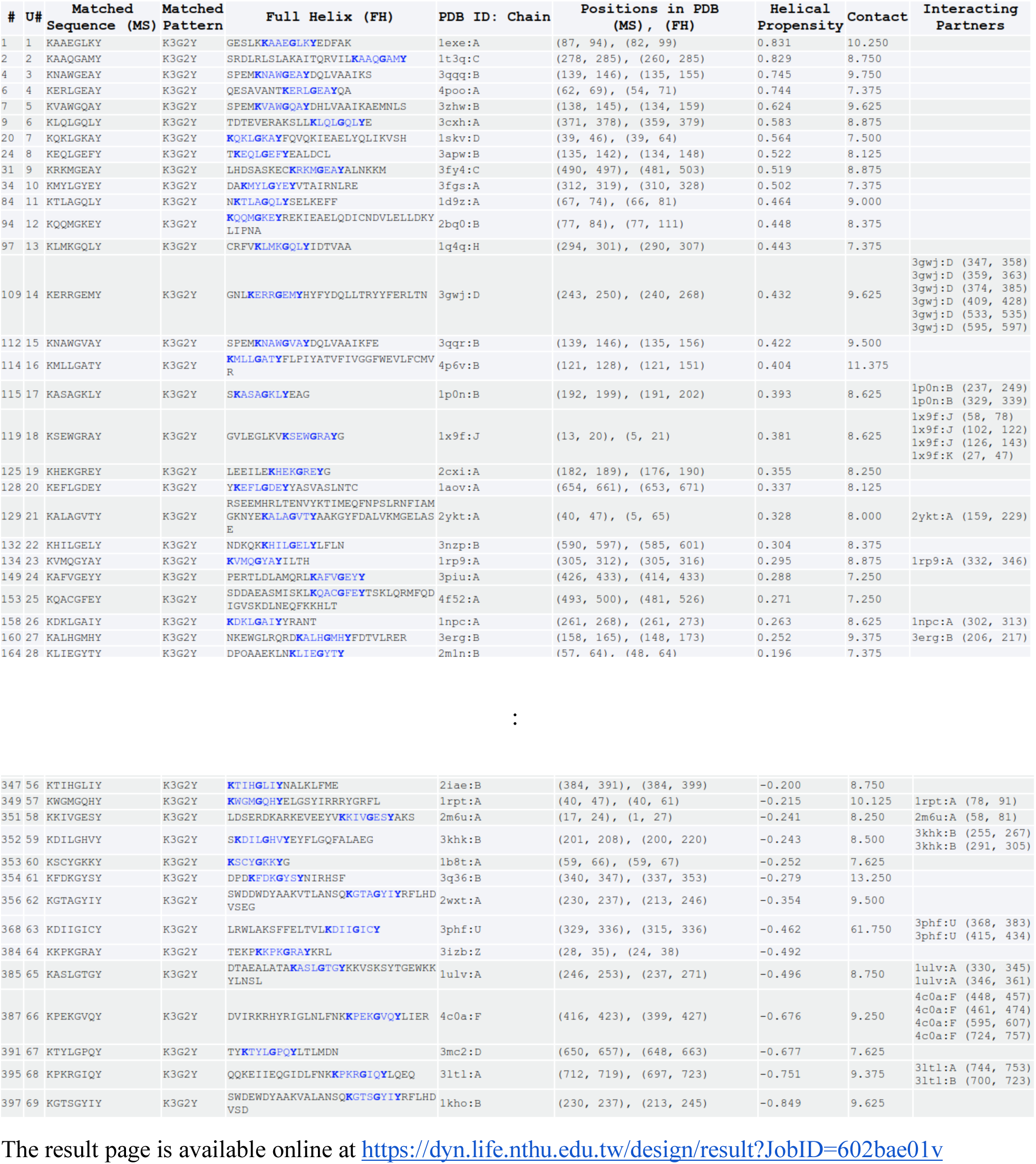
69 unique helical sequences obtained from the TP-DB matching the pattern K***G**Y given the query of [K 3 G 2 Y]

**Table S3.**
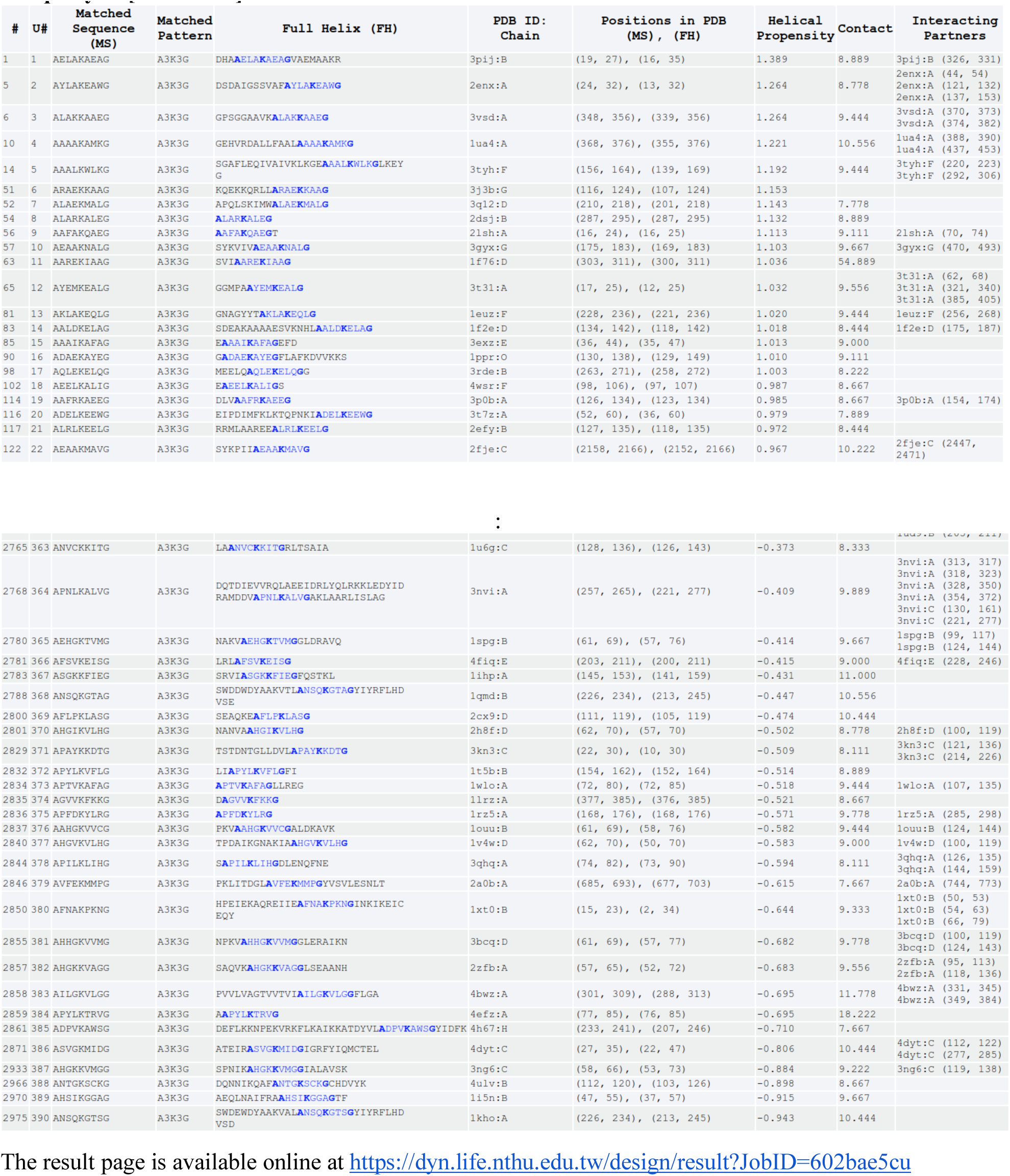
390 unique helical sequences obtained from the TP-DB for the pattern A***K***G given the query of [A 3 K 3 G]

**Table S4.**
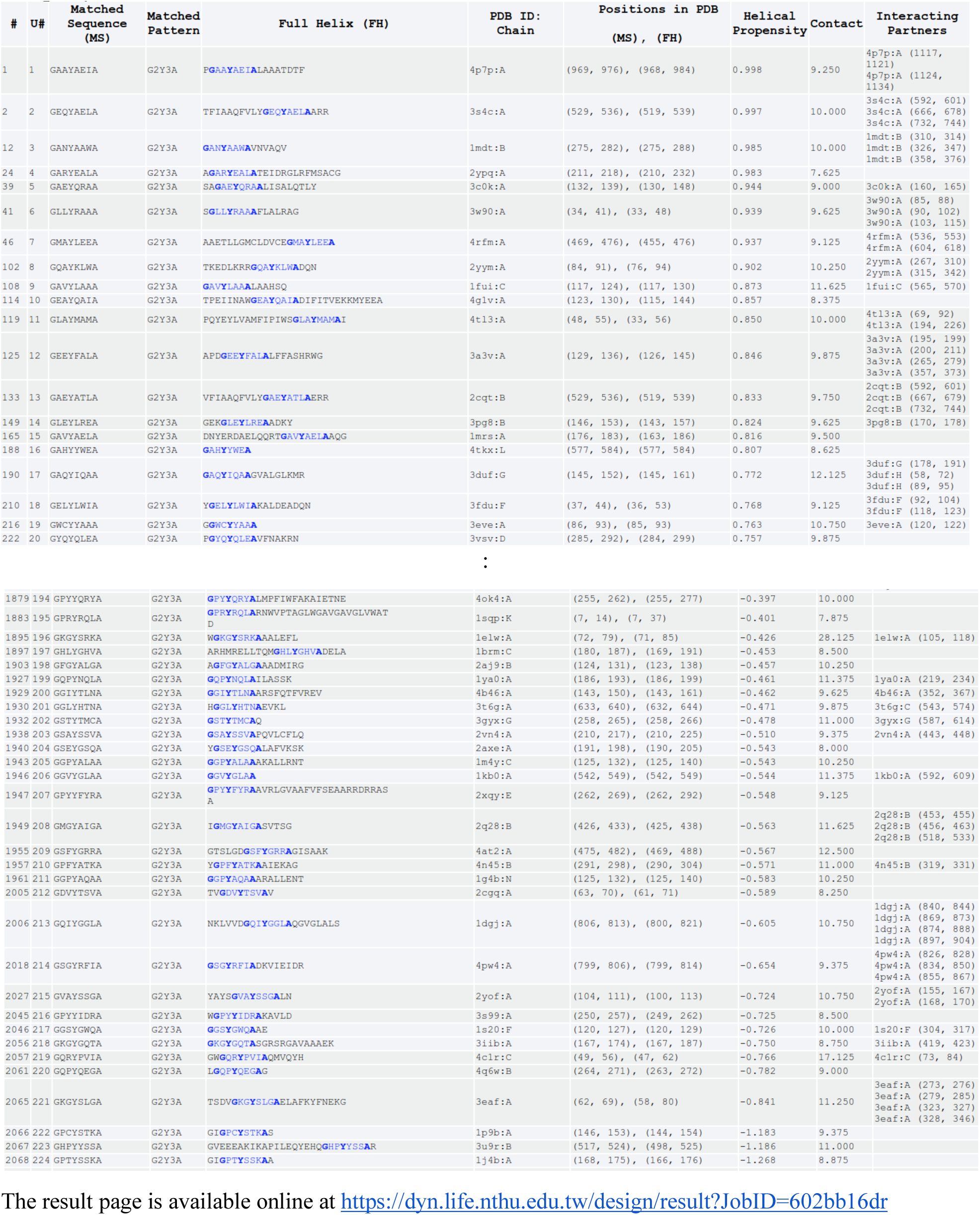
224 unique helical sequences obtained from the TP-DB for the pattern G**Y***A given the query of [G 2 Y 3 A]

**Table S5.**
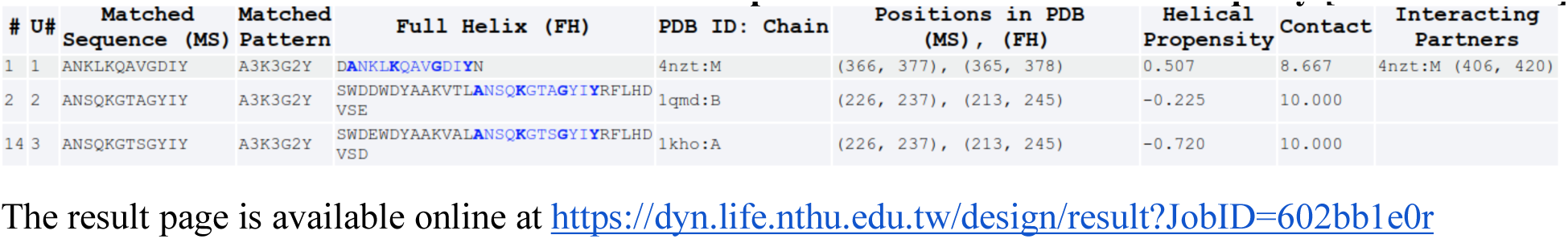
Results obtained from the TP-DB for pattern A***K***G**Y with query [A 3 K 3 G 2 Y]

**Table S6.**
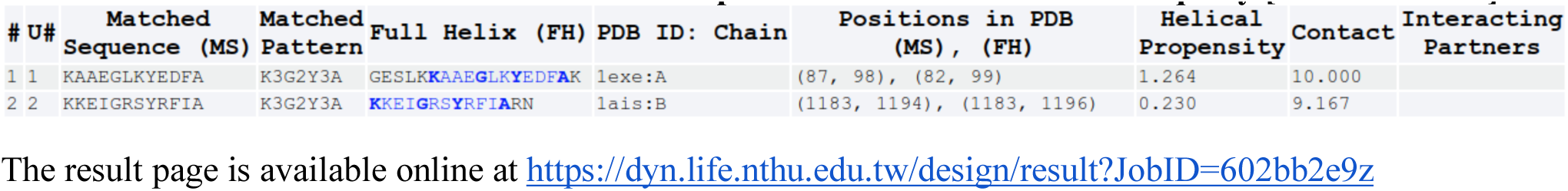
Results obtained from the TP-DB for pattern K***G**Y***A with query [K 3 G 2 Y 3 A]

**Table S7.**
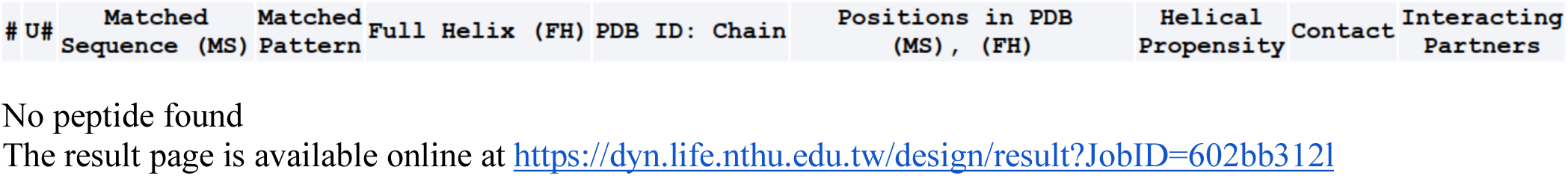
Results obtained from the TP-DB for pattern A***K***G**Y***A with query [A 3 K 3 G 2 Y 3 A]

**Table S8.**
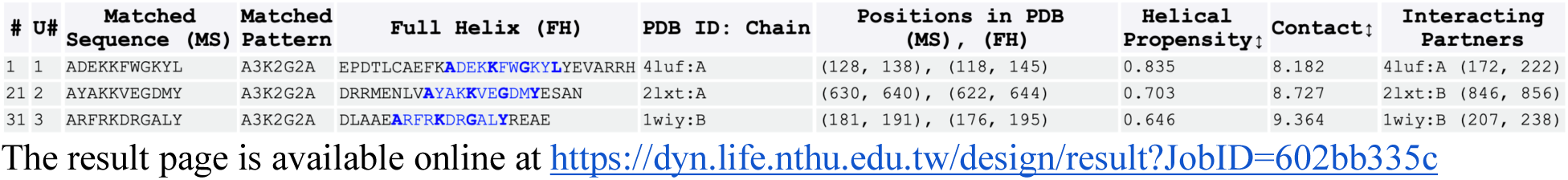
Results obtained from the TP-DB for pattern A**/***K**/***G**/***Y**/***A with query [A 2,3 K 2,3 G 2,3 Y 2,3 A]

We selected the helical PP2A-like peptides found from queries [A 3 K 3 G 2 Y] (**Table S5**), [K 3 G 2 Y 3 A] **(Table S6**), [A 2,3 K 2,3 G 2,3 Y 2,3 A] (**Table S8**), as well as the wide-type PP2A helix as control, for further investigations through all-atom MD-based binding free energy change (ΔG) evaluation by MM/PBSA (see Supporting Methods). The results (**Table S9**) showed that seven of the newly found helixes, with a binding ΔΔG < 0.0 kcal/mol, bound Sgo1 stronger than the wide-type PP2A did, suggesting a potential PPI blocking function for these peptides. Among them, the number one peptide “**A**YA**K**KVE**G**DM**Y**”, with a binding affinity ∼10 kcal/mol stronger than the wild-type peptide, can be a promising peptide subject to further experimental validation by isothermal calorimetry (ITC) assays and/ or NMR HSQC spectra. Peptide design is an elaborated miniature of protein design, and we expect that herein proposed methodologies can be applied for the design of therapeutic α-helical peptides for other medicinal purposes.

**Table S9.** Seven of the newly found helixes, with binding ΔΔG less than 0.0 kcal/mol, have the potentials of binding stronger to SGO1’s helix than the control

| Candidate Helix<br>(Matched Pattern) | Helix Source,<br>PDB ID: Chain | Helical<br>Propensity<br>Score | Contact<br>Number | Binding $\Delta\Delta G$<br>(kcal/mol) | In which Table<br>above* (U#) |
| --- | --- | --- | --- | --- | --- |
| **AYAKKVEGDMY**<br>(A 2,3 K 2,3 G 2,3 Y) | 2lxt:A | 0.703 | 8.545 | -10.11 ± 0.43 | **Table S8 (2')** |
| **KKEIGRSYRFIA**<br>(K 3 G 2 Y 3 A) | 1ais:B | 0.230 | 9.167 | -9.20 ± 0.32 | **Table S6 (2)** |
| **KAAEGLKYEDFA**<br>(K 3 G 2 Y 3 A) | 1exe:A | 1.264 | 10.000 | -8.04 ± 0.29 | **Table S6 (1)** |
| **AYAKKVEGDMYESA**<br>(A 2,3 K 2,3 G 2,3 Y 2,3 A) | 2lxt:A | 0.703 | 8.727 | -6.64 ± 0.45 | **Table S8 (2)** |
| **ANSQKGTAGYIY**<br>(A 3 K 3 G 2 Y) | 1qmd:B | -0.225 | 10.000 | -5.63 ± 0.33 | **Table S5 (2)** |
| **ARFRKDRGALYREA**<br>(A 2,3 K 2,3 G 2,3 Y 2,3 A) | 1wiy:B | 0.646 | 9.364 | -1.96 ± 0.46 | **Table S8 (3)** |
| **ARFRKDRGALY**<br>(A 2,3 K 2,3 G 2,3 Y) | 1wiy:B | 0.957 | 8.500 | -1.14 ± 0.38 | **Table S8 (3')** |
| **KTIHGLIY**<br>Control, from PP2A | 3fga:B | -0.200 | 9.250 | 0.00 |  |
| **ANSQKGTSGYIY**<br>(A 3 K 3 G 2 Y) | 1kho:A | -0.720 | 10.000 | 1.38 ± 0.39 | **Table S5 (3)** |
| **ADEKKFWGKYLY**<br>(A 2,3 K 2,3 G 2,3 Y) | 4luf:A | 0.957 | 8.500 | 4.31 ± 0.29 | **Table S8 (1')** |
| **ADEKKFWGKYLYEVA**<br>(A 2,3 K 2,3 G 2,3 Y 2,3 A) | 4luf:A | 0.835 | 8.182 | 4.66 ± 0.86 | **Table S8 (1)** |
| **ANKLKQAVGDIY**<br>(A 3 K 3 G 2 Y) | 4nzt:M | 0.507 | 8.667 | 5.11 ± 0.32 | **Table S5 (1)** |

\*The earlier Tables that listed the matched sequences are denoted at the rightmost column. The number in the parenthesis is the serial number for the unique (U#) sequence in the corresponding Table of interest. A prime following a U#, such as 2', indicates that the sequence in that row of this table is a subsequence of the original sequence presented in the earlier Tables.

## Supplementary Methods

### Production of recombinant Helicobactor pylori neutrophil-activating protein (HP-NAP) and maltose-binding protein (MBP)

Recombinant *H. pylori* neutrophil-activating protein (HP-NAP) was expressed in *E. coli* BL21(DE3) cells harboring the expression plasmid pET42a-NAP and purified by either two consecutive gel-filtration chromatography as previously described^60^ or a small-scale DEAE Sephadex negative mode batch chromatography as previously described^61^. Maltose-binding protein (MBP) was prepared the same as the procedure for production of MBP fused with the polypeptide containing residues Arg77 to Glu116 of HP-NAP as described below except that *E. coli* BL21(DE3) cells harboring the pMALc2g expression vector was used for expression.

### Cloning of HP-NAP_R77-E116_ into a MBP fusion protein expression vector

The plasmid DNA pET42a-NAP encoding a *napA* gene from *H. pylori* strain 26695 [GenBank:AE000543.1, Gene: HP0243] was prepared as previously described^60^. The DNA fragment coding for polypeptide containing residues Arg77 to Glu116 of HP-NAP (HP-NAP_R77-E116_), which contains the D-Y-K-x-x-[DE] motif, was amplified by PCR from the plasmid pET42a-NAP using the forward and reverse primers containing BamHI and HindIII site, respectively. The forward primer is 5’-ATAAGGATCCCGTGTTAAAGAAGAAACTAAAAC-3’ and the reversed primer is 5’-TTAATAAGCTTTAATTCTTTTTCAGCGGTGTTAGAG-3’. The PCR reaction was carried out with 10 ng plasmid DNA pET42a-NAP as a template and KAPA HiFi PCR Kit (Kapa Biosystems, Inc.) in a Mastercycler Gradient 5331 (Eppendorf, Germany). An initial denaturing phase of 95 °C for 5 min was followed by 39 cycles of 98 °C for 20 sec, 67 °C for 15 sec, and 72 °C for 15 sec. A final elongation phase of 72 °C for 2 min was also included. The amplified DNA fragments encoding HP-NAP_R77-E116_ were then cloned into pJET1.2/blunt vectors using the CloneJET PCR Cloning Kit (Thermo Fisher Scientific Inc.). The resulting plasmid was designated as pJET1.2/blunt-HP-NAP_R77-E116_. The insert was sequenced to confirm the correct DNA sequence. The correct insert was digested from pJET1.2/blunt-HP-NAP_R77-E116_ with BamHI and HindIII and then cloned into the pMALc2g expression vector^62^. The resulting plasmid was designated as pMALc2g-HP-NAP_R77-E116_.

### Production of MBP-tagged HP-NAP_R77-E116_

*E. coli* BL21(DE3) cells harboring pMALc2g-HP-NAPR77-E116 were streaked on a lysogeny broth (LB) agar plate containing 100 µg/ml ampicillin and incubated at 37 °C for 16 hr. A single colony was picked and inoculated into 4 ml of LB containing 100 µg/ml ampicillin and the culture was incubated at 37 °C with shaking at 170 rpm for 16 hr. A volume of 2 ml of the overnight culture was inoculated into 200 ml LB containing 100 µg/ml ampicillin and the inoculated culture was incubated at 37 °C with shaking at 170 rpm for 2 hr until the OD600 reached 0.5. The expression of MBP-tagged HP-NAPR77-E116 was induced by the addition of isopropyl β-D-1-thiogalactopyranoside (IPTG) to a final concentration of 0.3 mM and the culture was incubated at 37 °C with shaking at 180 rpm for 3 h until the OD600 reached 1.7. Then, the cells were centrifuged at 6,000 x g at 4 °C for 15 minutes to remove the supernatant and the cell pellets were stored at -70 °C.

The cell pellet from a 200 ml culture of *E. coli* expressing recombinant MBP-tagged HP-NAP_R77-E116_ were re-suspended in 20 ml of ice-cold buffer containing 20 mM Tris-HCl, pH 7.4, 200 mM NaCl, 1 mM ethylenediaminetetraacetic acid (EDTA), and 1 mM dithiothreitol (DTT), plus 0.1% (v/v) protease inhibitor mixture (PI mix). The PI mix contained 0.13 M phenylmethylsulfonyl fluoride (PMSF), 0.03 M N-alpha-tosyl-L-lysyl-chloromethyl ketone (TLCK), and 0.03 M N-tosyl-L-phenylalanyl-chloromethyl ketone (TPCK). The bacterial suspensions were disrupted by Emulsiflex C3 high-pressure homogenizer (Avestin) operated at a range of 15,000-20,000 psi for 7 times at 4 °C. The lysates were centrifuged at 30,000 x g at 4 °C for 1 hr to separate insoluble and soluble proteins by using a Hitachi himac CP80WX ultracentrifuge (Hitachi Koki Co. Ltd., Tokyo, Japan). Then, 5 mL supernatant containing the soluble proteins were loaded onto a 1-ml MBPTrap HP column (GE Healthcare Bio-Sciences), which was pre-equilibrated with 20 mM Tris-HCl, pH 7.4, 200 mM NaCl, 1 mM EDTA, and 1 mM DTT, at a flow rate of 0.5 ml/min at 4 °C by ÄKTA Purifier. The column was eluted with 20 mM Tris-HCl, pH 7.4, 200 mM NaCl, 1 mM EDTA, 1 mM DTT, and 10 mM maltose at a flow rate of 1 ml/min at 4 °C by ÄKTA Purifier. The flow-through and elution fractions were analyzed by SDS-PAGE on a 12% gel. The elution fractions containing recombinant MBP-tagged HP-NAPR77-E116 were collected and concentrated to a concentration higher than 1 mg/ml.

### Recombinant HP-NAP-based enzyme linked immunosorbent assay (ELISA)

Nunc MaxiSorp ninety-six-well enzyme linked immunosorbent assay (ELISA) plates (Nunc, Rochester, NY, USA) were coated with 0.3 µg of recombinant HP-NAP in 100 µl of bicarbonate buffer (pH 9.0) for each well at room temperature for 16 hr. Each well was washed with 300 µl of phosphate buffered saline (PBS), pH 7.4, containing 20 mM Na2HPO4, 1.47 mM KH2PO4, 137 mM NaCl, and 2.7 mM KCl, with the addition of 0.1% tween-20 (PBS-T) three times for 10 min each time. The wells were blocked with 250 µl PBS with 1% bovine serum albumin (BSA) for 2 hr and then washed with 300 µl of PBS-T buffer three times for 10 min each time. The anti-FLAG M2 antibody (Sigma-Aldrich, Cat# F-3165) and its corresponding mouse IgG antibody (Sigma-Aldrich, Cat# I5381) at a concentration of 600 ng/ml and the hybridoma culture supernatant containing mouse monoclonal antibody MAb 16F4^63^ against HP-NAP at a dilution of 1:5000 in 100 µl of PBS-T buffer containing 1% BSA were added into each well. The plate was incubated at room temperature for 1 hr and then the wells were washed with 300 µl of PBS-T buffer three times for 10 min each time. The horseradish peroxidase-conjugated goat anti-mouse secondary antibody (Jackson ImmunoResearch) at a dilution of 1:10000 in 100 µl of PBS-T buffer containing 1% BSA was loaded into each well. The plate was incubated at room temperature for 1 hr and then the wells were washed with 300 µl of PBS-T buffer three times for 10 min each time. The color was developed using 3,3’,5,5’-tetramethylbenzidine (TMB) peroxidase substrate (Thermo Scientific). The reaction was terminated by the addition of 2 N H_2_SO_4_, and the absorbance at 450 nm was measured by Bio-rad iMark microplate absorbance reader (Hercules, CA).

### Western blot analysis

Western blotting was performed essentially the same as previously described^64^. The membrane was probed with either anti-FLAG M2 antibody (Sigma-Aldrich) at a concentration of 1 µg/ml or the hybridoma culture supernatant containing mouse monoclonal antibody MAb 16F4^63^ against HP-NAP at a dilution of 1:2000.

### Finding PP2A-like Peptides

Knowing that the K374, G378, and Y381 of PP2A plays important roles in PP2A’s favourable interactions with Sgo1 (see **Fig S4**), we set out to search for a peptide with a similar amino acid pattern and which may interact better than with Sgo1 and thereby out compete/displace PP2A and bind to Sgo1. Therefore, we searched the developed TP-DB for the following queries: **[K 3 G 2 Y]**, **[A 3 K 3 G], [G 2 Y 3 A], [A 3 K 3 G 2 Y], [K 3 G 2 Y 3 A], [A 3 K 3 G 2 Y 3 A], and [A 2,3 K 2,3 G 2,3 Y 2,3 A].** The resulting peptides from each query are ranked based on their helical propensity score and contact number.

### Minimization and MD simulations of Martini Coarse-grained (CG) models in vacuum and assessing the stability of complexes

To search for PP2A-like helical peptides to block the interaction between Sgo1 and PP2A, 69 sequences (https://dyn.life.nthu.edu.tw/design/result?JobID=602bae01v) that matched the pattern “K***G**Y” in PP2A helix (see **Fig S4**; the three anchoring residues of PP2A helix **K**_374_**T**_375_**I**_376_**H**_377_**G**_378_**L**_379_**I**_380_**Y**_381_ that interacts with Sgo1 are K374, G378 and Y381) were chosen for affinity assessment. Methodologically, it was equivalent to computationally modify (mutate) the five spacing residues T375, I376, H377, L379 and I380 in PP2A helix “K<u>TIH</u>G<u>LI</u>Y” into corresponding residues in every of these 69 sequences. In other words, we only replaced the alpha-helical segment “K_374_TIHGLIY_381_” in PP2A [PDB:3FGA; chain B] by every of the 69 helical stretches that matched the pattern “K***G**Y” and the segment was shown to interact with the helix “V_75_KEAQDIILQLRKECYYL_92_” in Sgo1 [PDB:3FGA; chain D]. Subsequently, all 69 complexes comprising the Sgo1 helix and mutated PP2A helix were converted into coarse-grained (CG) models by the web service CHARMM-GUI^65^. Both terminals of the helical peptides in all complexes were set to be electrically neutral, and 69 CG models with PP2A mutants as well as the counterpart of wide-type were energy-minimized and then briefly equilibrated for 1 ps at 20 fs time step by MD simulations running on GROMACS package^66^ with MARTINI force field^67, 68^. The pressure and temperature were maintained at 1 bar and 310K, respectively. The cutoff distances for both Coulomb and Van der Waal were set to be 12Å.

MD simulations were conducted in vacuum with modest positional restraints (spring constant = 2.5 kcal/mol/Å^2^) exerted on the backbone beads for 1 ps at 20 fs time step, and the potential energies at the final step were recorded for the analysis to obtain the binding stability between the targeted helix in Sgo1 and mutated (or wild-type) PP2A segments.

Besides the top-ranked peptides by Martini forcefield from the aforementioned 69 complexes, we also searched potential helical binders from TP-DB using at least 4 anchoring residues in the search pattern (**Tables S5** to **S8**). We carried out all-atom molecular dynamics simulations for these peptides to assess their interactions with the targeted helix of Sgo1 using MM/PBSA^69, 70^ (see below). The top 11 identified helices (**Table S9**), searched from the queries [A 3 K 3 G 2 Y], [K 3 G 2 Y 3 A] and [A 2,3 K 2,3 G 2,3 Y 2,3 A], were first superimposed onto the template helix of the PP2A [PDB: 3FGA; chain B; residue 372 to 383] with their common residues K374, G378, and Y381 (as shown in Fig S3), before the simulations.

We carried out the all-atom molecular dynamics simulations by AMBER16^71, 72^ package, using ff14SB forcefield^73^ for protein and ionsjc_tip3p for ions^74^ in explicit solvent. Each molecular system to be simulated was placed in a periodic box and solvated with TIP3P water. All the input files for MD simulations were prepared using tLeap^71^ from AmberTools16.

Energy minimization was performed in three stages - first, with weak harmonic positional restraints (spring constant = 0.5 kcal/mol/Å^2^) on all atoms except for the water/solvent atoms; second, with weak harmonic restraints on the CA atoms of amino acids; lastly, without restraints. Each of the first two stages of energy minimization is composed of 5,000 steps, with the first 2500 steps (for each energy minimization stage) carried out using steepest descent algorithm and the remaining 2,500 steps carried out using conjugate gradient algorithm.

Following the energy minimization, each simulation system was slowly heated to 310K while applying weak harmonic positional restraints on the CA atoms. Each system was then equilibrated at the temperature reached without any positional restraints prior to production MD simulations. Each production MD simulation ran for 500 ns at 2 fs time step. Non-bonded interactions were evaluated up to a cut-off distance of 10 Å where it was switched off with a cubic spline switch function. Particle Mesh Ewald method^23^ was used to calculate full electrostatic interactions energies. All temperature regulations were done using Langevin thermostat (with a collision frequency, γ, of 2 ps^-1^) and all pressure control was done with Berendsen barostat^75^.

Binding free energy change (binding ΔG) was calculated from the MD simulations trajectories using AMBER16’s implementation of Molecular Mechanics/Poisson-Boltzmann Surface Area (MM/PBSA)^69, 70^.

