## Supplementary Results for "Secondary Structure Motifs Made Searchable to Facilitate the Functional Peptide Design"

***for***

*^1^Institute of Bioinformatics and Structural Biology, National Tsing-Hua University, Hsinchu 30013, Taiwan; ^2^Graduate Institute of Medical Genomics and Proteomics, National Taiwan University College of Medicine, Taipei 10055, Taiwan; ^3^Bioinformatics Program, Institute of Information Sciences, Academia Sinica, Taipei 11529, Taiwan;* *^4^Department of Computational and Systems Biology, School of Medicine, University of Pittsburgh, Pittsburgh, PA, 15213, USA; ^5^Institute of Molecular and Cellular Biology, National Tsing Hua University, Hsinchu 30013, Taiwan;* *^6^Praexisio Taiwan Inc. New Taipei 22180, Taiwan; ^7^Department of Life Science, National Tsing Hua University, Hsinchu 30013, Taiwan*; *^8^Physics Division, National Center for Theoretical Sciences, Hsinchu 30013, Taiwan.*

¶ Authors share equal contributions

**Supplementary Results**

*Chemical structures of POPC and POPG lipids*

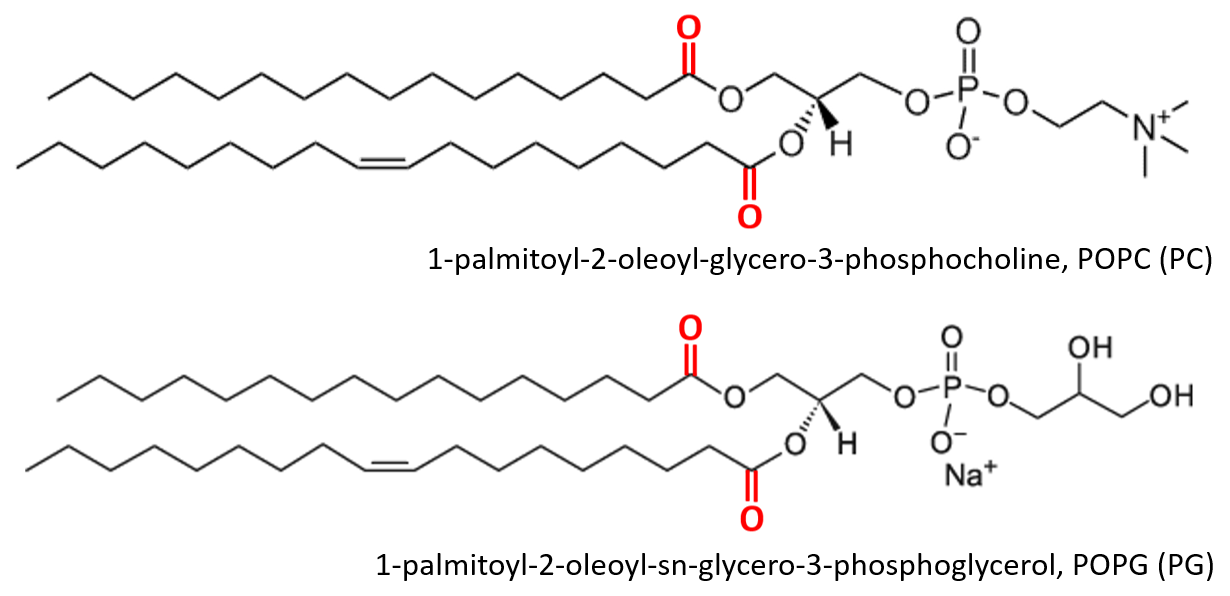

**Figure. S1** Chemical structure of POPC (PC) and POPG (PG). All images modified from Avanti® product page (Product ID: 840457 and 850457). The carbonyl oxygen in the lipids are highlighted in red, and these oxygen atoms are represented as red points in Fig. 3A. The atoms of the inserted AMPs below these oxygen atoms are considered to interact with the aliphatic tails of the lipids.

*Natural insertion of three AMPs into bacteria membrane*

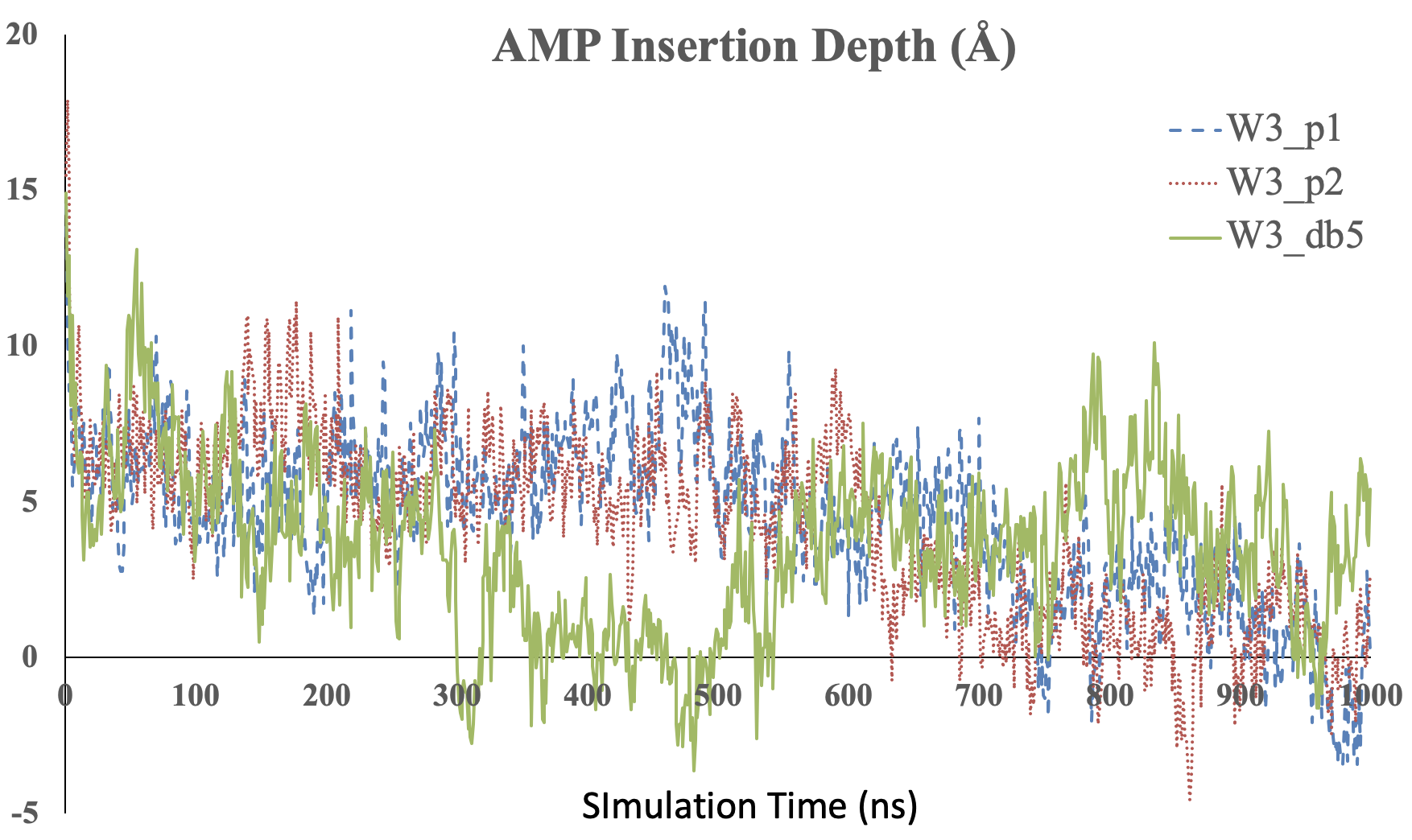

**Figure S2.** The relative position of the center of mass (COM) of AMPs to the COM of phosphor atoms in the upper leaflet consisting of 40 POPC/POPG (3:1) lipid molecules, mimicking bacteria membrane, is plotted against the simulation time. All the MD simulations were conducted for 1 microsecond (us) at body temperature and normal pressure (1 atm) using OpenMM^20^ with CHARMM36 forcefield^21,22^.

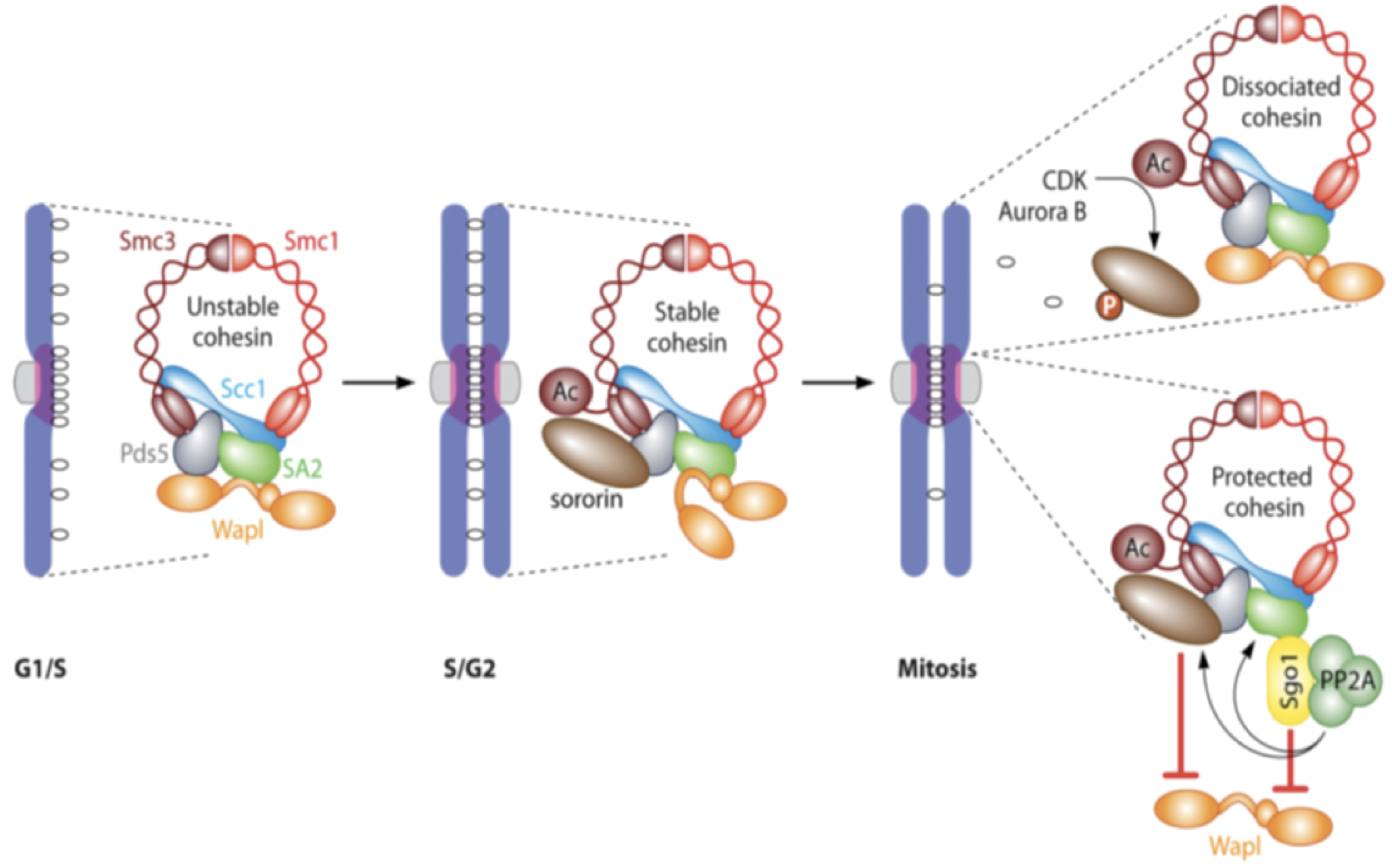
**Figure S3** The overview of Sgo1-PP2A protecting centromeric cohesin. During mitosis, phosphorylation of SA2 and Sororin cause cohesin to dissociate from chromosome arm. Cdk1-mediated phospho-Sgo1 recruits PP2A at centromere to counteract phosphorylation of SA2 and Sororin and maintains centromeric cohesin to tether sister chromatids together. Image is adopted from Figure 3 in Marston’s review article^59^.

We noted that the complex of Sgo1-PP2A has been solved by x-ray crystallography (PDB ID: 3FGA) and the two molecules bind with each other through an helix-helix interface (one helix from each molecule)^11^. To design anticancer peptides as blocker against Sgo1-PP2A interaction, the helical stretch in PP2A (residues from 352-392 in chain B of PDB 3FGA), known to form the helix-helix interaction with another helix in Sgo1 (residues from 75 to 92 in chain D of PDB 3FGA) (Fig S4), is chosen as the candidate target for further investigation. Evidenced from the x-ray structure, the helix-helix interaction is visibly apparent to be the most important essential interacting elements at the protein-protein interface. As shown in Figure S3a, K374, G378 and Y381 in the helical stretch “KTIHGLIY” could be the anchoring residues on PP2A to mediate the Sgo1-PP2A interaction, which is supported by our calculation of energetics using *in silico* alanine scanning (Fig S4b).

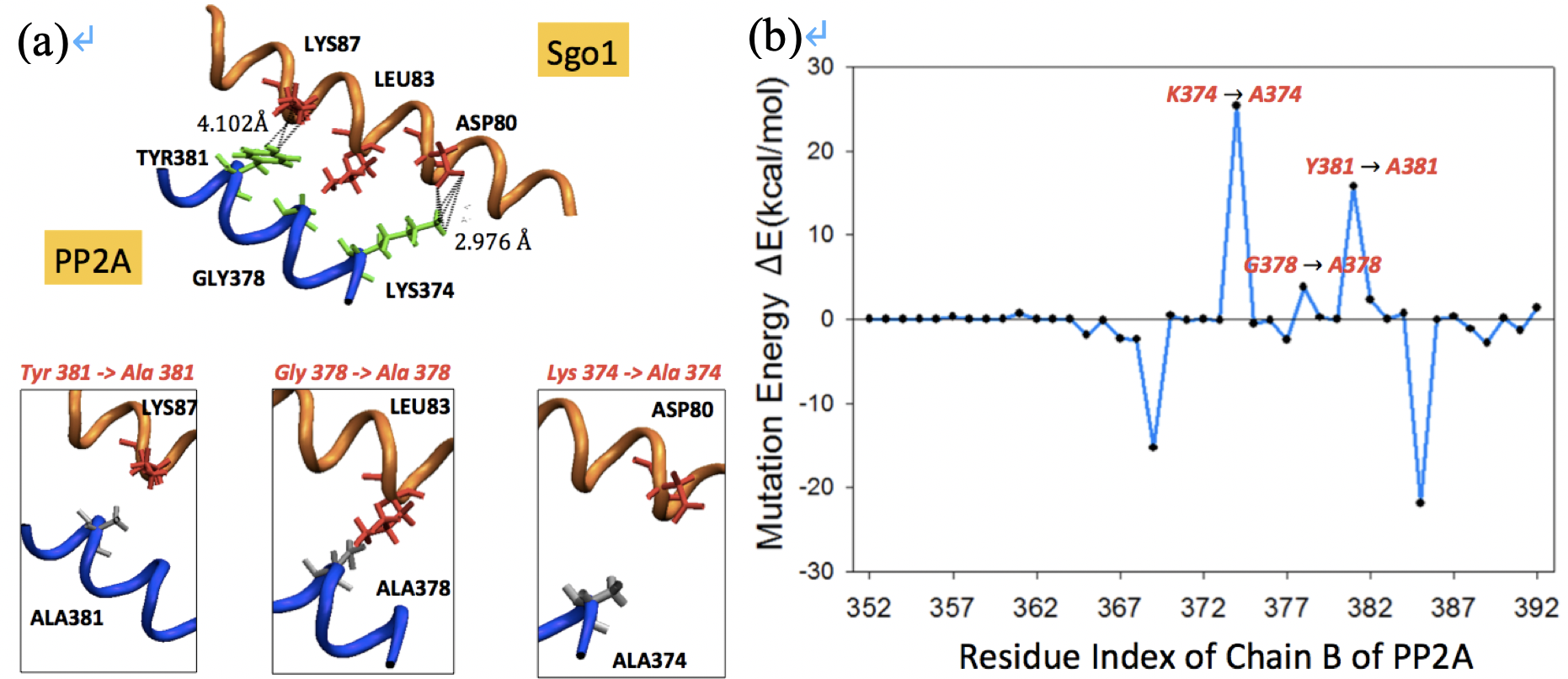

**Figure S4** Protein-protein interaction (PPI) between Sgo1 and PP2A through helix-helix interactions. Panel **(a, upper)** shows only the PPI interface between Sgo1 and PP2A, while the rest of proteins (3FGA; Xu et al., 2009) are hidden for clarity. Sgo1-PP2A interaction is mediated by two helices, one from Sgo1 (PDB: 3FGA; V_75_KEAQDIILQLRKECYYL_92_ in chain D) and the other from PP2A (PDB: 3FGA; K_369_THMNKTIHGLIYNALK_385_ in chain B) . Computational alanine scanning for every residue in the PP2A helical fragment (from residue 352 to 392 of chain B) is carried out, and in panel (**b**) energetic contribution for each residue mutated into alanine is assessed (from residue 369 to 385 of chain B) where a large positive value (e.g. for K374A, G378A and Y381A) indicates the importance of the residues in binding, and a large negative value suggests mutation into alanine favors the binding. The non-bonded energy is evaluated by NAMD package (Phillips et al., 2005) using CHARMM 36 forcefield (Huang and MacKerell, 2013). Here in panel **(b)**, the mutation energy for the n-th residue, ∆En, is defined as ∆En = Yn - X0 where X0 is the non-bonded energy of original complex between Sgo1 and PP2A (for the 352-392 fragment), and Yn is the energy of the complex with the *n*-th residue in PP2A being mutated into alanine, after a short energy minimization.

To block the protein-protein interaction (PPI) between Sgo1 and PP2A, one could potentially modify part of the most essential interacting element in PP2A into a peptide that interacts with Sgo1 better than its template. As shown in Figure S4b, computational alanine scanning helps detecting a stretch from 370 to 384 in PP2A (chain B) dominantly contributing the affinity in the PPI of Sgo1-PP2A complex, especially from K374, G378 and Y381 (i.e. aforementioned essential interacting element in PP2A). We then used this helical stretch, or the extended one with its flanking region from residue 364 to 388, including two energetically favored point mutations, K369A and K385A, as the starting template.

To find the helical peptides that bind the interface helix of Sgo1 stronger than PP2A does, we search the TP-DB for PP2A-like helical peptides using the queries summarized in Table S1.

As detailed in Supporting Methods, we first used the pattern “**K***G**Y**” to search TP-DB and discover 69 unique PP2A-like helical peptides (Table S1 and S2); we then carried out molecular dynamics simulations (MD) to assess how these peptides interact with Sgo1 at the helix-helix interface. Helices H60, H54, H61, H33 and H29 (**Fig S5**) are energetically promising for further experimental confirmation. Additionally, we examined other sets of peptides that met the patterns consisting of three anchoring residues (“A***K***G” and “G**Y***A”) or consisting of at least four residues (“A***K***G**Y”, “K***G**Y***A”, “A***K***G**Y***A” and “A**/***K**/***G**/***Y**/***A”**)** in TP-DB. All the corresponding results obtained from the queries are illustrated in **Table S3** to **S8**. Among them only the “A***K***G**Y***A” pattern (according to Figure S3b) returns no result from TP-DB.

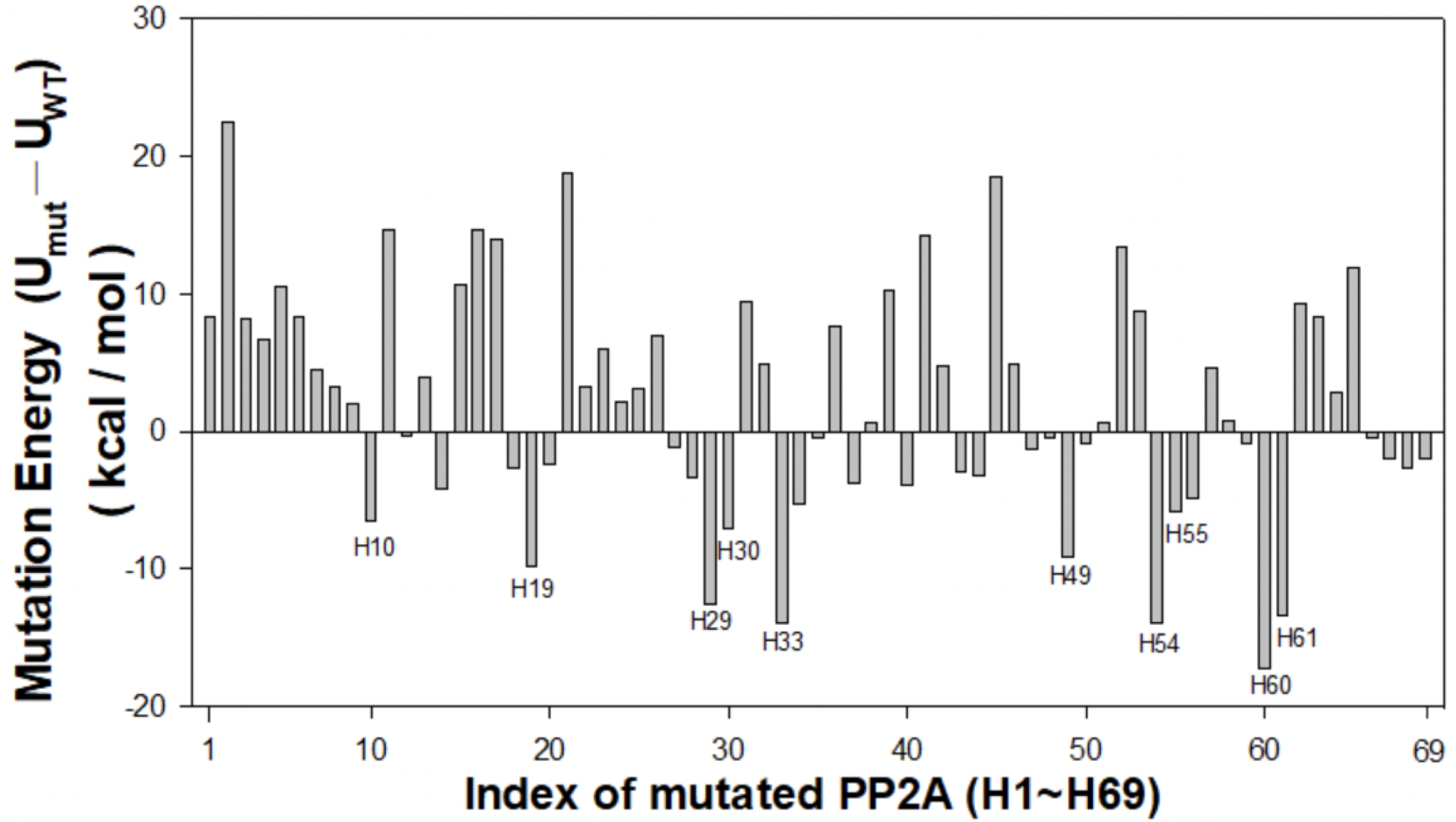

**Figure S5.** **Interactions between Sgo1 helix and PP2A-like peptides.** U_mut_ is the interaction energy between targeted Sgo1 helix and each of the 69 PP2A-like peptides (from H1 to H69) that match the pattern “K3G2Y”. In detail, the energy is calculated by replacing the residue T375, I376, H377, L379 and I380 in the “**K_374_**TIH**G**LI**Y_381_**” stretch of PP2A with the corresponding residues in each of the 69 pattern-matched (K3G2Y) helical peptides returned from TP-DB. U_WT_ is the interaction energy between Sgo1 helix and PP2A. Larger difference of interaction energies (U_mut_ - U_WT_) suggest stronger binding of PP2A-like peptides with the Sgo1 helix than that between PP2A and the Sgo1 helix. Top ten values are further labeled in the graph.

**Table S1**. PP2A-like patterns searched against the developed helical peptide database

| Pattern | Query | # of Peptides Found | # of Unique Peptides Found |
| --- | --- | --- | --- |
| K *** G ** Y | K 3 G 2 Y | 398 | 69 |
| A *** K *** G | A 3 K 3 G | 2981 | 391 |
| G ** Y *** A | G 2 Y 3 A | 2079 | 225 |
| A *** K *** G ** Y | A 3 K 3 G 2 Y | 15 | 3 |
| K *** G ** Y *** A | K 3 G 2 Y 3 A | 3 | 2 |
| A *** K *** G ** Y *** A | A 3 K 3 G 2 Y 3 A | 0 | 0 |
| A**/***K**/***G**/***Y**/***A | A 2,3 K 2,3 G 2,3 Y 2,3 A | 34 | 3 |

**Table S2. 69 unique helical sequences obtained from the TP-DB matching the pattern K***G**Y given the query of [K 3 G 2 Y]**

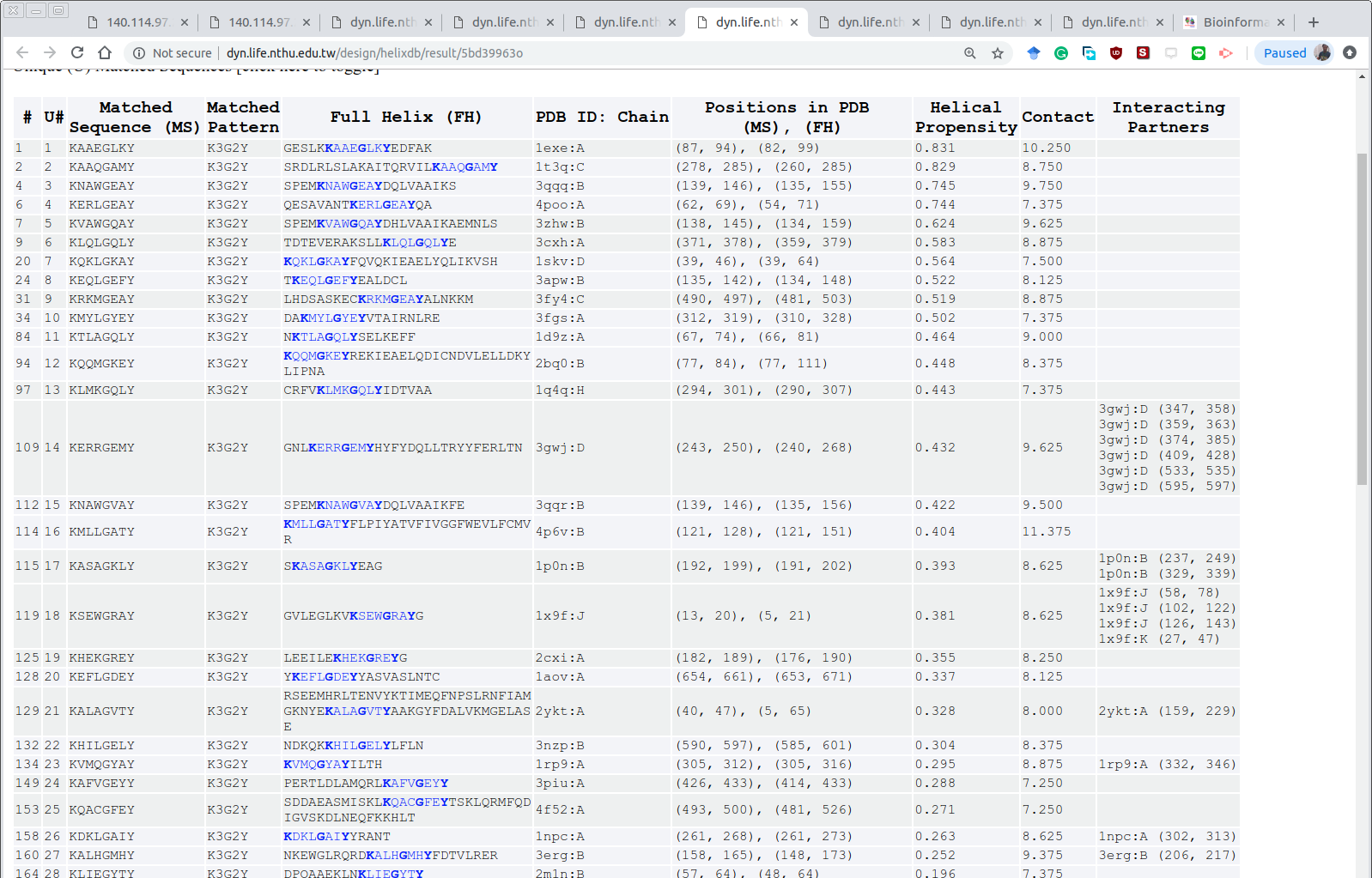

:

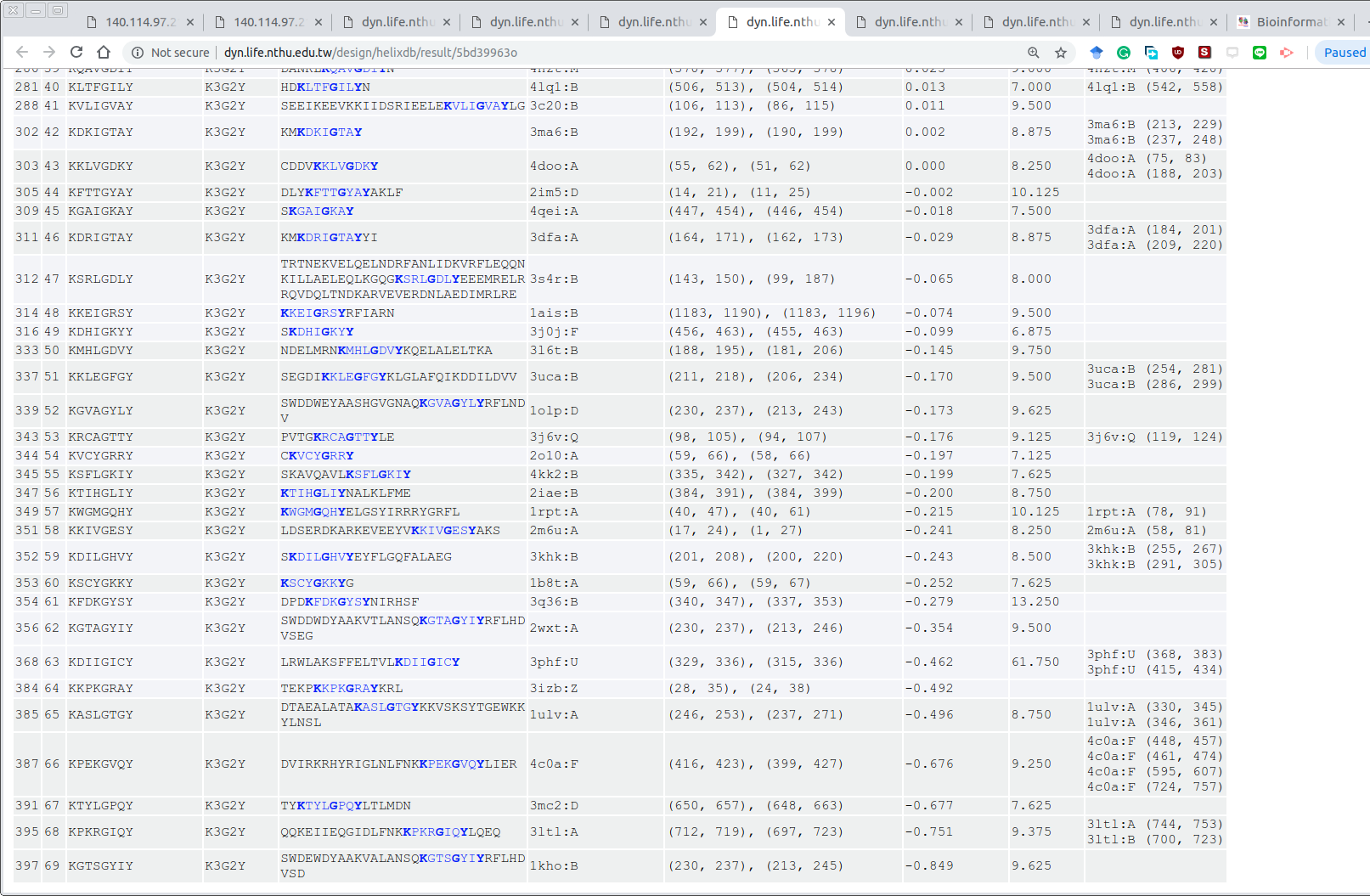

The result page is available online at <https://dyn.life.nthu.edu.tw/design/result?JobID=602bae01v>

**Table S3. 390 unique helical sequences obtained from the TP-DB for the pattern A***K***G given the query of [A 3 K 3 G]**

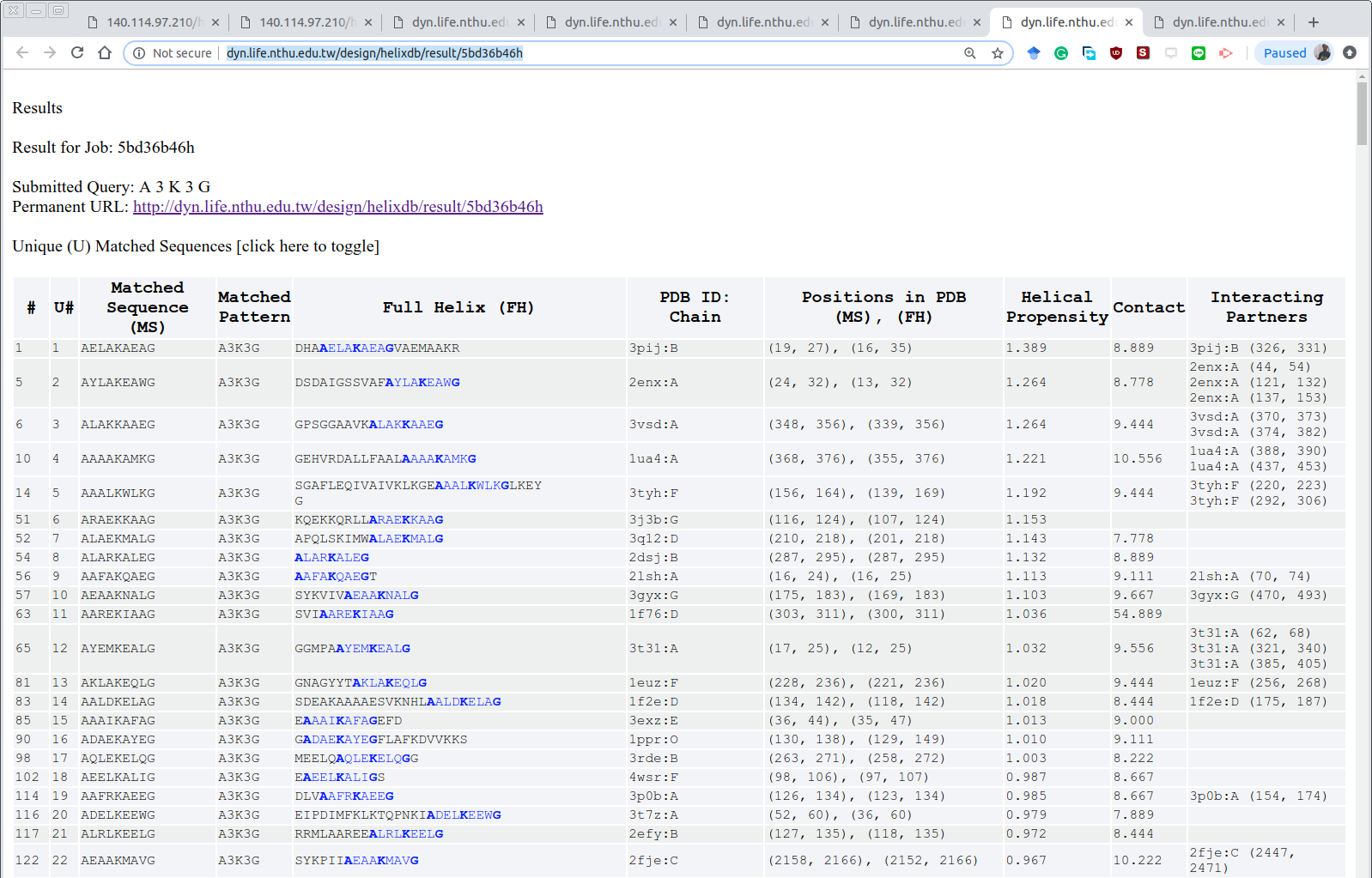

:

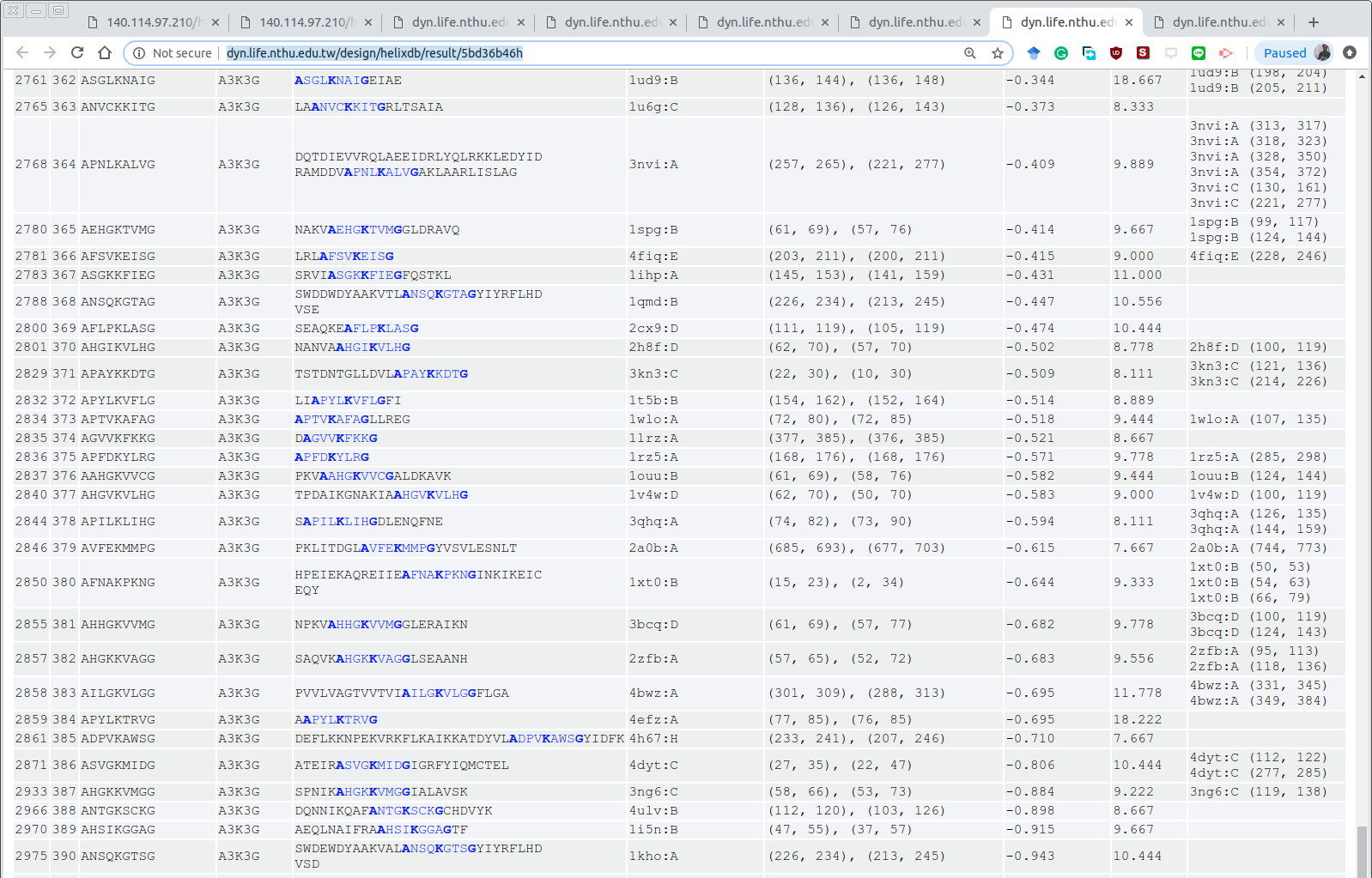

The result page is available online at <https://dyn.life.nthu.edu.tw/design/result?JobID=602bae5cu>

**Table S4. 224 unique helical sequences obtained from the TP-DB for the pattern G**Y***A given the query of [G 2 Y 3 A]**

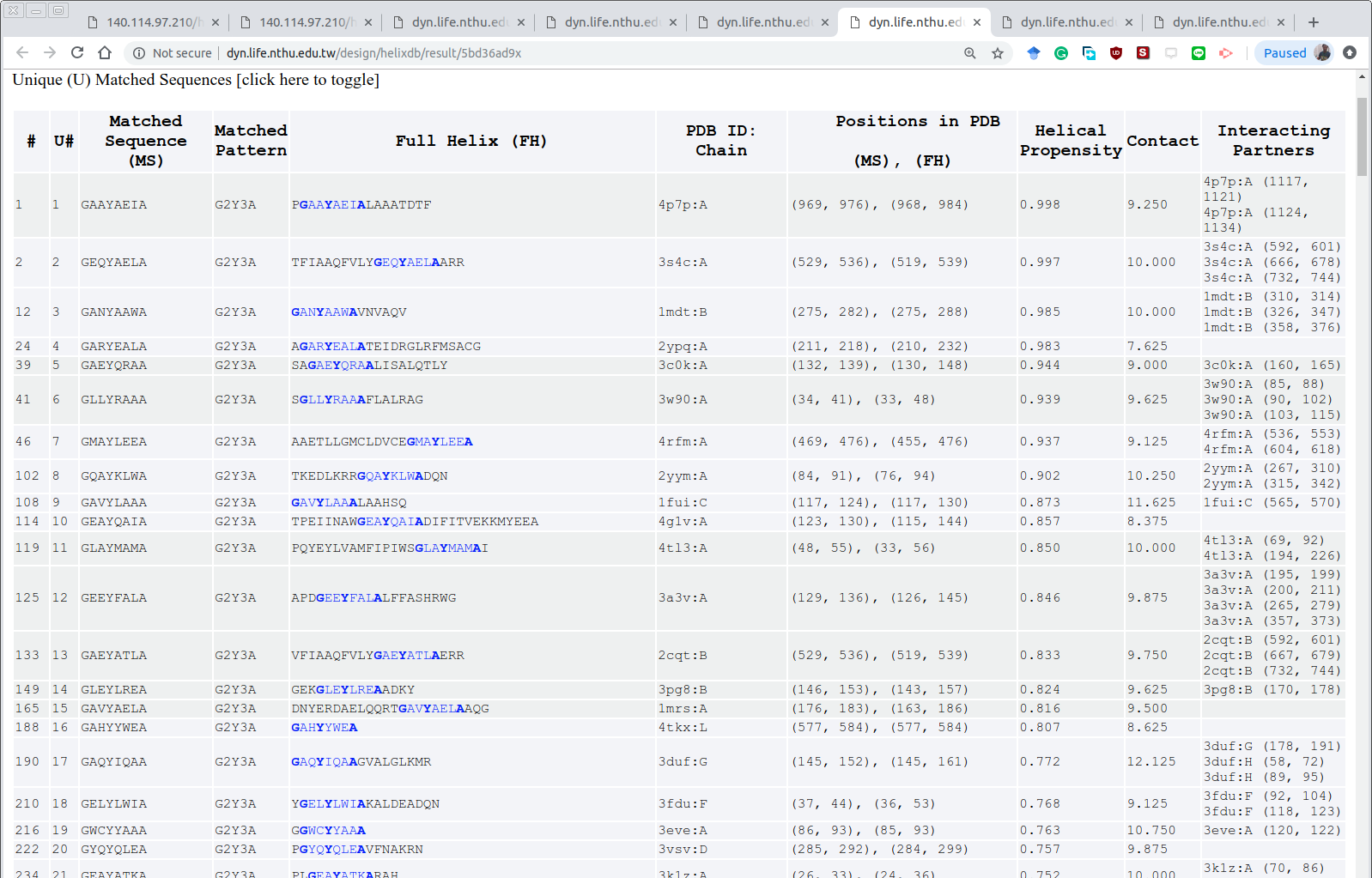
:

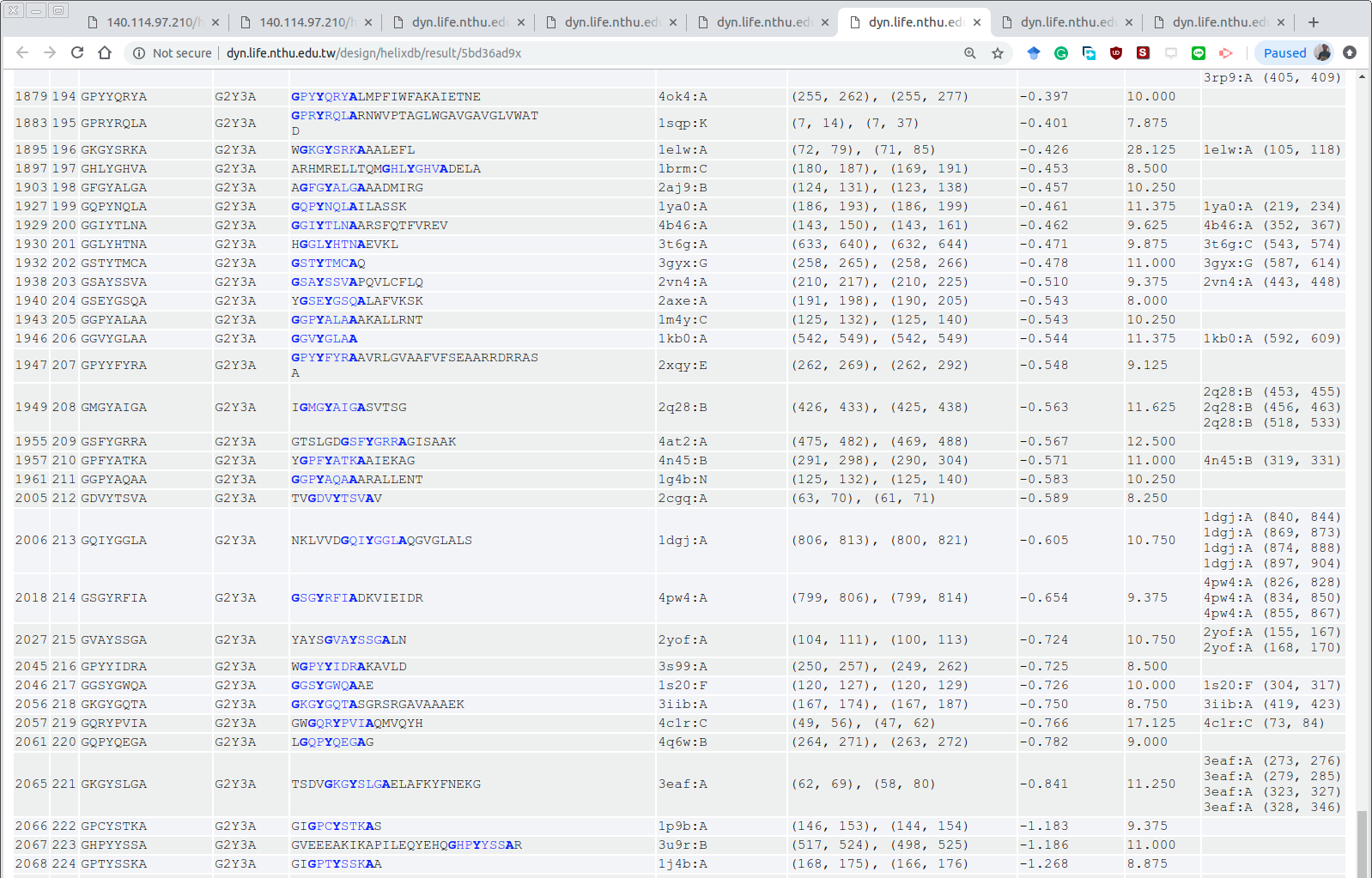

The result page is available online at <https://dyn.life.nthu.edu.tw/design/result?JobID=602bb16dr>

**Table S5. Results obtained from the TP-DB for pattern A***K***G**Y with query [A 3 K 3 G 2 Y]**

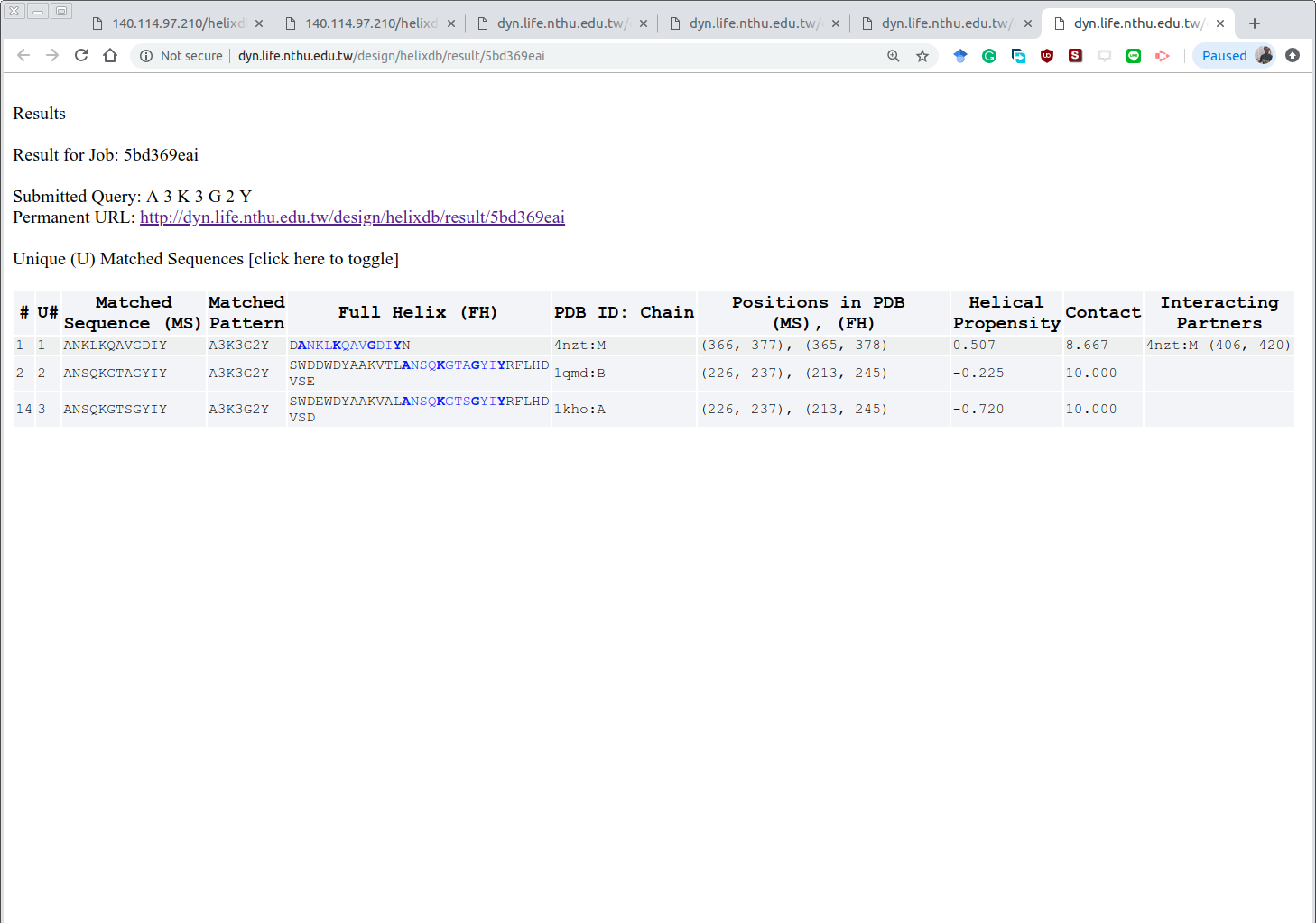

The result page is available online at <https://dyn.life.nthu.edu.tw/design/result?JobID=602bb1e0r>

**Table S6. Results obtained from the TP-DB for pattern K***G**Y***A with query [K 3 G 2 Y 3 A]**

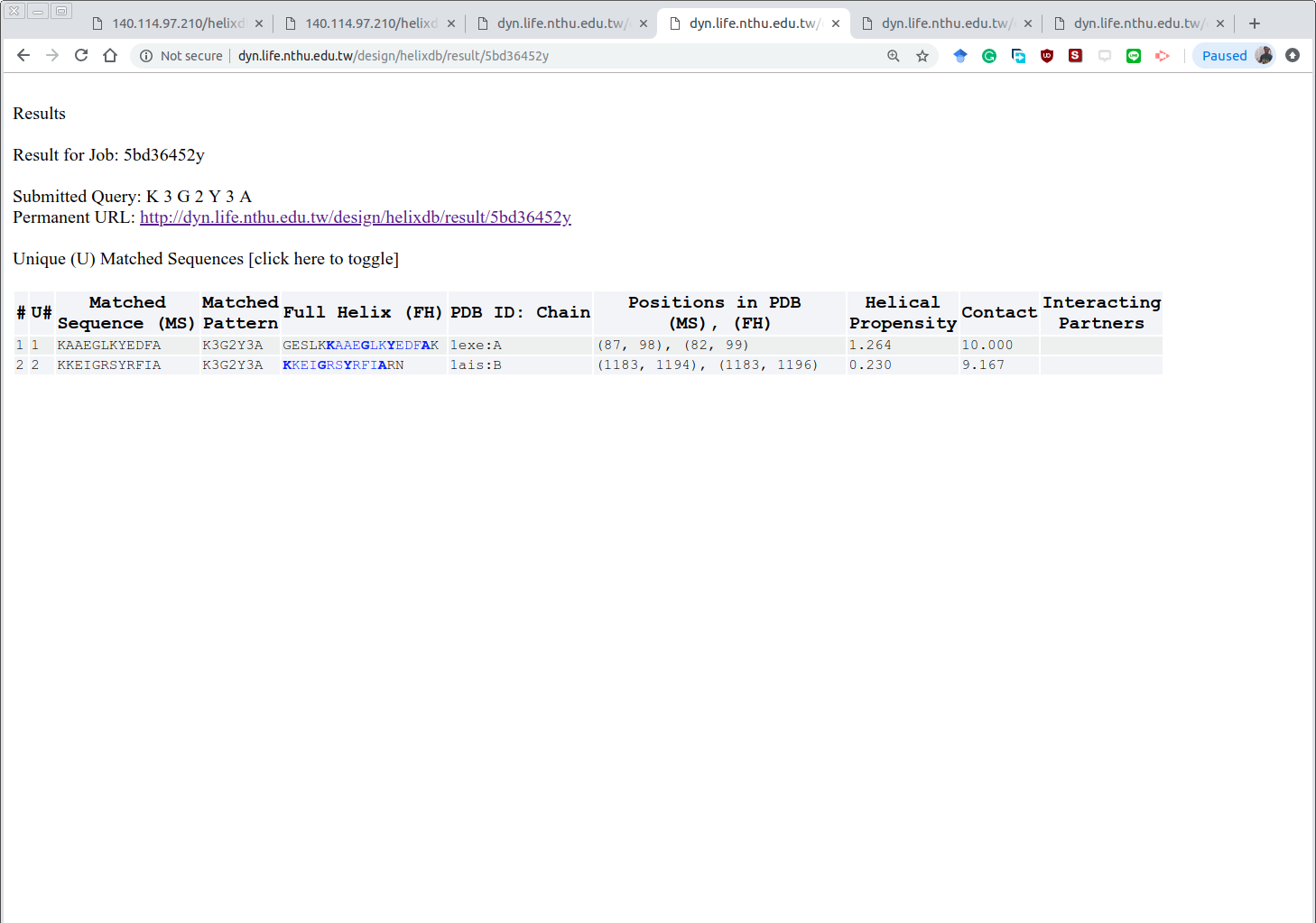

The result page is available online at <https://dyn.life.nthu.edu.tw/design/result?JobID=602bb2e9z>

**Table S7. Results obtained from the TP-DB for pattern A***K***G**Y***A with query [A 3 K 3 G 2 Y 3 A]**

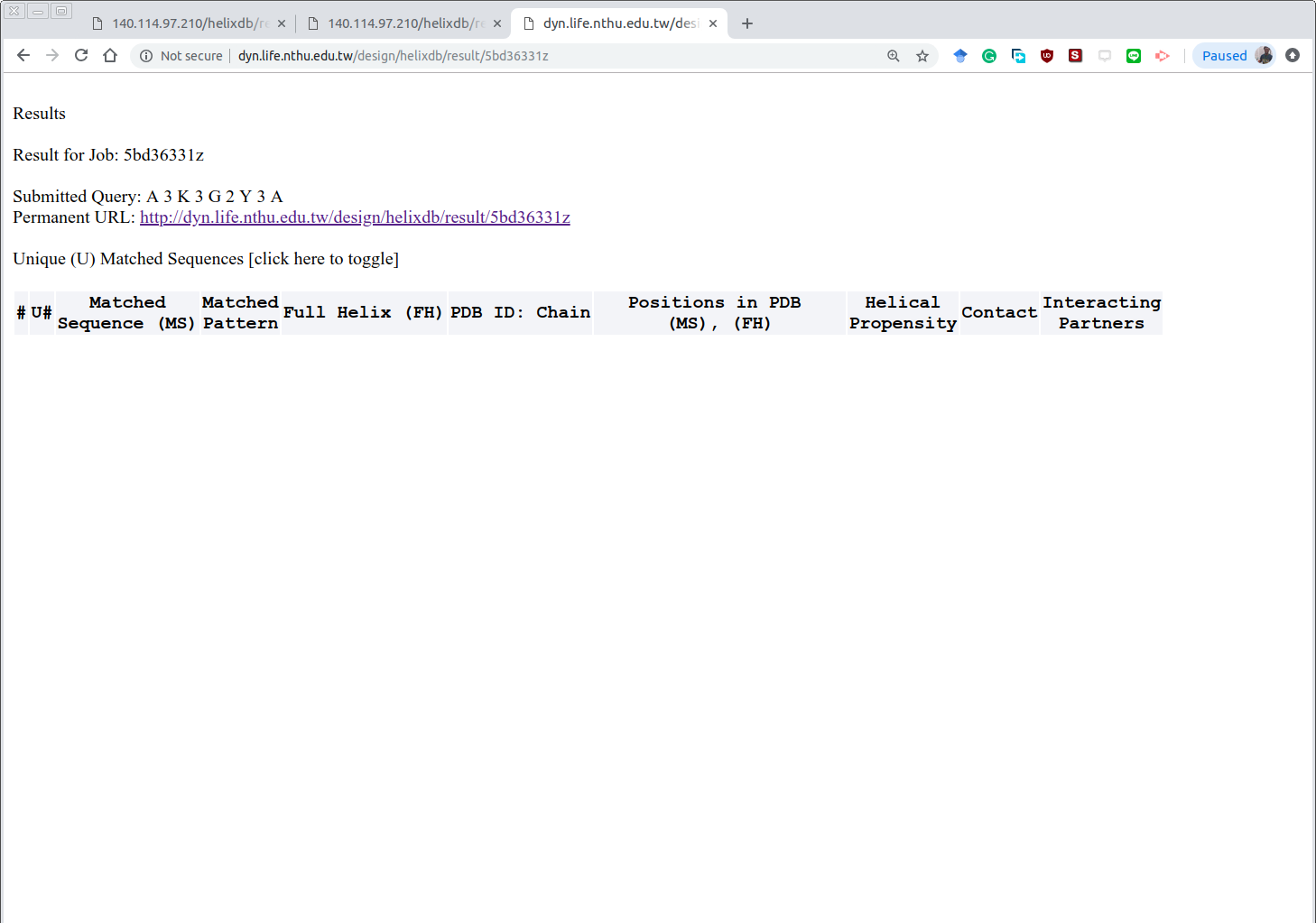

No peptide found

The result page is available online at <https://dyn.life.nthu.edu.tw/design/result?JobID=602bb312l>

**Table S8. Results obtained from the TP-DB for pattern** **A**/***K**/***G**/***Y**/***A with query [A 2,3 K 2,3 G 2,3 Y 2,3 A]**

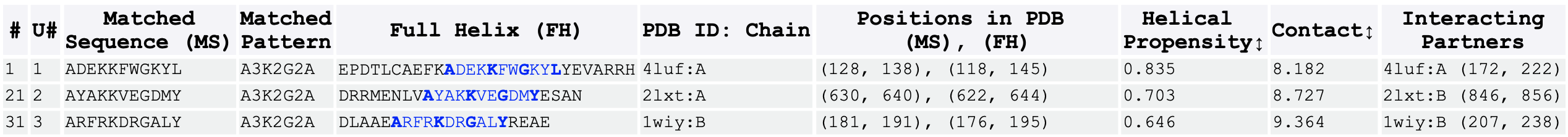

The result page is available online at <https://dyn.life.nthu.edu.tw/design/result?JobID=602bb335c>

We selected the helical PP2A-like peptides found from queries [A 3 K 3 G 2 Y] (**Table S5**), [K 3 G 2 Y 3 A] **(Table** **S6**), [A 2,3 K 2,3 G 2,3 Y 2,3 A] (**Table S8**), as well as the wide-type PP2A helix as control, for further investigations through all-atom MD-based binding free energy change (∆G) evaluation by MM/PBSA (see Supporting Methods). The results (**Table S9**) showed that seven of the newly found helixes, with a binding ∆∆G < 0.0 kcal/mol, bound Sgo1 stronger than the wide-type PP2A did, suggesting a potential PPI blocking function for these peptides. Among them, the number one peptide “**A**YA**K**KVE**G**DM**Y**”, with a binding affinity ~10 kcal/mol stronger than the wild-type peptide, can be a promising peptide subject to further experimental validation by isothermal calorimetry (ITC) assays and/ or NMR HSQC spectra. Peptide design is an elaborated miniature of protein design, and we expect that herein proposed methodologies can be applied for the design of therapeutic α-helical peptides for other medicinal purposes.

**Table S9. Seven of the newly found helixes, with binding ∆∆G less than 0.0 kcal/mol, have the potentials of binding stronger to SGO1's helix than the control**

| Candidate Helix  (Matched Pattern) | Helix Source, PDB ID: Chain | Helical Propensity Score | Contact  Number | Binding ∆∆G (kcal/mol) | In which Table above* (U#) |
| --- | --- | --- | --- | --- | --- |
| **A**YA**K**KVE**G**DM**Y**  (**A 2,3 K 2,3 G 2,3 Y**) | 2lxt:A | 0.703 | 8.545 | -10.11 ± 0.43 | **Table S8** (2') |
| **K**KEI**G**RS**Y**RFI**A**  (**K 3 G 2 Y 3 A**) | 1ais:B | 0.230 | 9.167 | -9.20 ± 0.32 | **Table S6** (2) |
| **K**AAE**G**LK**Y**EDF**A**  (**K 3 G 2 Y 3 A**) | 1exe:A | 1.264 | 10.000 | -8.04 ± 0.29 | **Table S6** (1) |
| **A**YA**K**KVE**G**DM**Y**ES**A**  (**A 2,3 K 2,3 G 2,3 Y 2,3 A**) | 2lxt:A | 0.703 | 8.727 | -6.64 ± 0.45 | **Table S8** (2) |
| **A**NSQ**K**GTA**G**YI**Y**  (**A 3 K 3 G 2 Y**) | 1qmd:B | -0.225 | 10.000 | -5.63 ± 0.33 | **Table S5** (2) |
| **A**RFR**K**DR**G**AL**Y**RE**A**  (**A 2,3 K 2,3 G 2,3 Y 2,3 A**) | 1wiy:B | 0.646 | 9.364 | -1.96 ± 0.46 | **Table S8** (3) |
| **A**RFR**K**DR**G**AL**Y**  (**A 2,3 K 2,3 G 2,3 Y**) | 1wiy:B | 0.957 | 8.500 | -1.14 ± 0.38 | **Table S8** (3') |
| **K**TIH**G**LI**Y**  **Control, from PP2A** | 3fga:B | -0.200 | 9.250 | 0.00 |  |
| **A**NSQ**K**GTS**G**YI**Y**  (**A 3 K 3 G 2 Y**) | 1kho:A | -0.720 | 10.000 | 1.38 ± 0.39 | **Table S5** (3) |
| **A**DEK**K**FW**G**KYL**Y**  (**A 2,3 K 2,3 G 2,3 Y**) | 4luf:A | 0.957 | 8.500 | 4.31 ± 0.29 | **Table S8** (1') |
| **A**DEK**K**FW**G**KYL**Y**EV**A**  (**A 2,3 K 2,3 G 2,3 Y 2,3 A**) | 4luf:A | 0.835 | 8.182 | 4.66 ± 0.86 | **Table S8** (1) |
| **A**NKL**K**QAV**G**DI**Y**  (**A 3 K 3 G 2 Y**) | 4nzt:M | 0.507 | 8.667 | 5.11 ± 0.32 | **Table S5** (1) |

*The earlier Tables that listed the matched sequences are denoted at the rightmost column. The number in the parenthesis is the serial number for the unique (U#) sequence in the corresponding Table of interest. A prime following a U#, such as 2', indicates that the sequence in that row of this table is a subsequence of the original sequence presented in the earlier Tables.
